# Visualization of functional and effective connectivity underlying auditory descriptive naming

**DOI:** 10.1101/2025.03.18.644036

**Authors:** Yu Kitazawa, Kazuki Sakakura, Hiroshi Uda, Naoto Kuroda, Riyo Ueda, Ethan Firestone, Min-Hee Lee, Jeong-Won Jeong, Masaki Sonoda, Shin-ichiro Osawa, Kazushi Ukishiro, Makoto Ishida, Kazuo Kakinuma, Shoko Ota, Yutaro Takayama, Keiya Iijima, Toshimune Kambara, Hidenori Endo, Kyoko Suzuki, Nobukazu Nakasato, Masaki Iwasaki, Eishi Asano

## Abstract

**Objective:** We visualized functional and effective connectivity within specific white matter networks in response to auditory descriptive questions.

**Methods:** We investigated 40 Japanese-speaking patients with focal epilepsy and estimated connectivity measures using cortical high-gamma dynamics and MRI tractography.

**Results:** Hearing a *wh*-interrogative at question onset enhanced inter-hemispheric functional connectivity, with left-to-right callosal facilitatory flows between the superior-temporal gyri, contrasted by functional connectivity diminution with right-to-left callosal suppressive flows between dorsolateral prefrontal regions. Processing verbs associated with concrete objects or adverbs increased left intra-hemispheric connectivity, with bidirectional facilitatory flows through extensive white matter pathways. Questions beginning with *what*, compared to *where*, induced greater neural engagement in the left posterior inferior-frontal gyrus at question offset, linked to enhanced functional connectivity and bidirectional facilitatory flows to the temporal lobe neocortex via the arcuate fasciculus. During overt responses, inter-hemispheric functional connectivity was enhanced, with bidirectional callosal flows between Rolandic areas, and individuals with higher IQ scores exhibited less prolonged neural engagement in the left posterior middle frontal gyrus.

**Conclusions:** Visualization of directional neural interactions within white matter networks during overt naming is feasible.

**Significance:** Phrase order may influence network dynamics in listeners, even when presented with auditory descriptive questions conveying similar meanings.

## 1. INTRODUCTION

Human communication fundamentally involves listening and answering. Existing neurobiological models explain the locations and timing of the cerebral cortex areas that manage language and speech processes (Poeppel et al., 2012; Price, 2012; Hagoort and Indefrey, 2014; Bornkessel-Schlesewsky et al., 2015; Chang et al., 2015; Tremblay and Dick, 2016; Skeide and Friederici, 2016; Binder, 2017; Pulvermüller, 2023). The essential processes include the initial processing of acoustic sounds by the auditory cortex in the left superior temporal gyrus (STG), which are then transformed into spoken sounds through neural communications with the left superior and middle temporal gyri (MTG) (Fontolan et al., 2014; Binder, 2017; Brefczynski-Lewis and Lewis, 2017; Nakai et al., 2017; Fridriksson et al., 2018; Forseth et al., 2020; Hamilton et al., 2021; Lu et al., 2021; Rocchi et al., 2021; Bhaya-Grossman and Chang, 2022; Nourski et al., 2022; Leonard et al., 2024). The extensive temporal lobe neocortices, including the left middle and inferior temporal gyri (ITG), are involved in assigning meaning to these vocal elements as words and sentences (Binder and Desai, 2011; Hamberger et al., 2014; Nakai et al., 2017; Forseth et al., 2018; Fridriksson et al., 2018; Rolls et al., 2022; Coletta et al., 2024). The left frontal lobe, including the left posterior inferior frontal gyrus (pIFG), collaborates with the temporal lobe neocortices to formulate an internal response to a heard utterance, and the arcuate fasciculus is suggested to play a major role in information transfer between these structures (Petersson et al., 2012; Hagoort and Indefrey, 2014; Gajardo-Vidal et al., 2021; Rocchi et al., 2021; Rolls et al., 2022; Coletta et al., 2024). This mental representation of the verbal response is transferred to the left precentral gyrus, which is responsible for initiating the physical act of speaking (Flinker et al., 2015; Nakai et al., 2017; Lu et al., 2021; Coletta et al., 2024). A classic model infers that the neural information is unidirectionally transferred from the left temporal to the left frontal lobe (Geschwind, 1970). Contemporary models propose bidirectional interactions between specific pairs of cortical regions (Poeppel et al., 2012; Chang et al., 2015; Flinker et al., 2015; Tremblay and Dick, 2016; McClelland et al., 2020; Lorca-Puls et al., 2021; Wang et al., 2021; Rolls et al., 2022; Pulvermüller, 2023; Fitz et al., 2024); for instance, neural interaction is suggested to occur between the left temporal and frontal lobes, playing a pivotal role in the semantic and syntactic processing required to comprehend a spoken sentence and in the conversion of perceived auditory sounds into corresponding mouth movements for speech production (Poeppel et al., 2012; Cogan et al., 2014; Bornkessel-Schlesewsky et al., 2015; Rolls et al., 2022). Modern models also propose the enhancement and diminution of specific regions and networks, optimizing the processing required at given moments (Snyder et al., 2010; Geranmayeh et al., 2014; Nakai et al., 2017; Kucewicz et al., 2019; Pulvermüller, 2023; Fitz et al., 2024; Tuckute et al., 2024). Many models also propose an information exchange between the left and right cerebral hemispheres (Geschwind, 1970; Poeppel et al., 2012; Skeide and Friederici, 2016). Despite advancements, current models have not visualized the direction and strength of neural information flows through specific white matter pathways on the order of hundreds of milliseconds in a language task involving overt speech. To address these limitations, our study aims to develop a novel functional brain atlas that serves as a comprehensive neurobiological model of auditory naming. This atlas is designed to visualize the neural information flows through white matter pathways, capturing their dynamics during distinct moments of auditory naming processing.

To generate this functional brain atlas, we first determined the timing and locations of high-gamma amplitude modulation (70-110 Hz) on intracranial EEG (iEEG) recording during a language task.

This task involved listening to a question and responding verbally. The measurement of language task-related high-gamma augmentation is a method used in many tertiary epilepsy surgery centers for functional brain mapping. This technique has been validated by electrical stimulation mapping and by analyzing postoperative cognitive outcomes (Arya et al., 2018; Sonoda et al., 2022). Our previous iEEG study of 65 English-speaking patients with drug-resistant focal epilepsy demonstrated that postoperative language impairment was predicted by the resection of sites showing high-gamma augmentation elicited by naming to questions beginning with a *wh-* interrogative (Sonoda et al., 2022). *Wh*-questions are commonly and naturally used in everyday language or conversational contexts, and the advantage of using *wh-*questions as auditory stimuli includes the minimal task instruction needed for participants. We have found that both children and adults readily understand our auditory descriptive naming task and intuitively respond to *wh-* questions (Nakai et al., 2017). High-gamma activity serves as an outstanding summary measure of neural activation with its amplitude tightly correlated to the neural firing rate on single neuron recordings (Ray et al., 2008; Rich and Wallis, 2017; Leszczyński et al., 2020), hemodynamic responses on functional MRI (fMRI) (Nir et al., 2007; Kunii et al., 2013b), and glucose metabolism on positron emission tomography (Nishida et al., 2008). In the current study, we visualized the dynamics of functional connectivity modulations between pairs of distinct cortical regions via specific white matter. We declared that functional connectivity was significantly enhanced (or diminished) if two cortical regions showed simultaneous, significant, and sustained high-gamma augmentation (or attenuation) and also if these cortical regions were directly connected by a white matter tract on tractography (Kitazawa et al., 2023; Ono et al., 2023; Ueda et al., 2024). We then employed transfer entropy-based effective connectivity analysis (Ito et al., 2011; Firestone et al., 2023), to quantify the intensity and direction of neural information flow through each white matter tract that showed significant functional connectivity modulation during a given movement. We declared that a facilitatory (or suppressive) information flow occurred from one cortical region to another if, across such a white matter tract, an increase (or decrease) in high-gamma amplitude was predictive of a subsequent increase (or decrease) in a distinct cortical region. By employing the dynamic tractography imaging technique (Sonoda et al., 2021; Kitazawa et al., 2023; Sakakura et al., 2023; Ueda et al., 2024), we animated temporal changes in the neural information flows along given white matter pathways.

We designed connectivity analyses to address the inherent limitation of spatial sampling in iEEG by leveraging group-level high-gamma measures, an approach similarly adopted in various forms by other investigators. Common methodologies include constructing a virtual brain model, wherein neural propagation is inferred from the timing and amplitude of high-gamma activity recorded at anatomically matched electrode sites across multiple patients. Consequently, data from different patient groups collectively contribute to the iEEG recordings at specific anatomical locations. For instance, Kunii et al. (2013a) inferred neural propagation using high-gamma amplitudes recorded from 1,512 electrode sites across 21 patients performing various language tasks. Similarly, Avanzini et al. (2016) inferred propagation patterns of somatosensory information based on group-level high-gamma activity elicited by median nerve stimulation, utilizing data from 11,983 cortical electrode sites across 99 patients. Burke et al. (2014) grouped spatially aligned electrodes from 98 patients, visualizing high-gamma augmentation sequences during memory tasks and migration of neural activation from posterior perceptual regions to medial and ventrolateral temporal areas. Solomon et al. (2017) conducted a comprehensive analysis of inter-regional communication during a memory task using 30,064 electrode sites from 294 patients, examining connectivity across 1,458 ROI pairs both within and across hemispheres through aggregated patient data from different groups of patients. Individual-level iEEG connectivity analyses remain valuable but are inherently limited to examining neural communication within narrowly defined cortical regions due to restricted spatial sampling. Even extensive studies involving tens of thousands of electrodes acknowledge these constraints (Solomon et al., 2017), recognizing that individual analyses alone cannot comprehensively assess inter-and intra-hemispheric connectivity, as no single patient has sufficiently broad cortical coverage. Given that electrode placement is dictated solely by clinical necessity, it is ethically and practically infeasible to expect comprehensive bilateral sampling of homotopic nonepileptic regions in patients undergoing invasive evaluation for drug-resistant focal epilepsy. Thus, group-level approaches help overcome these sampling limitations and provide insights into normative neural connectivity patterns, including interhemispheric neural communication.

Some readers may question whether combining iEEG signals recorded from different patient groups into a virtual brain is valid for connectivity analysis. Our position is “yes”; group-level analysis is a feasible and reliable method to estimate neural communication shared by the general population. When computing group-level summary measures (e.g., mean values), increasing the number of electrode sites minimizes the influence of individual patient variability, enhances the signal-to-noise ratio at each region of interest (ROI), and improves the generalizability of connectivity findings. Exclusion of electrode sites affected by epileptiform discharges will minimize the risk of pathological activity contaminating the connectivity measures. Separate analysis of iEEG signals aligned to stimulus and response alleviate the effects of variance in response time within and across patients on connectivity measures. In our recent iEEG study (Kanno et al., 2025), we visualized intrahemispheric and interhemispheric functional connectivity at the whole-brain level during a naming task using data from 7,792 non-epileptic electrode sites sampled across 106 English-speaking patients. In that study, we measured high gamma-based functional connectivity using 50-ms time windows sliding every 10 ms and found that cortices exhibiting increased functional connectivity at specific times relative to stimulus or response had a higher likelihood of naming-related symptoms elicited by electrical stimulation mapping. Specifically, functional connectivity measured at 60 ms after stimulus offset showed the strongest association with stimulation-induced receptive aphasia (Spearman’s rho = +0.54, p-value: 2.5 × 10⁻⁵), whereas connectivity measured 380 ms before response onset was most strongly associated with stimulation-induced expressive aphasia (Spearman’s rho: +0.78; p-value: 3.9 × 10^-12^) (Kanno et al., 2025). These findings support our position that iEEG-based connectivity modulations can be reliably estimated by aggregating high-gamma activity recorded across multiple populations, even using brief time windows.

In the present study, participants were native Japanese-speaking patients with drug-resistant focal epilepsy (see the detailed demographics of patients in **Table S1**). They were assigned an auditory descriptive naming task during extraoperative iEEG recording, wherein they needed to overtly answer three-phrase questions with two distinct structures. One type of question began with a *wh*-interrogative followed by a combination of two concrete phrases, while the others started with a combination of two concrete phrases followed by a *wh*-interrogative. These structures are commonly used by Japanese-speaking individuals, and the phrase order in Japanese is known for its flexibility (Saito, 1992). Conversely, in English, a *wh*-question typically begins with a *wh*-interrogative. Therefore, we anticipated that our iEEG study of Japanese-speaking individuals would provide insights into the network dynamics related to phrase order. We also expected that effective connectivity analysis of iEEG would reveal the directions of neural information flows between cortical pairs showing functional connectivity enhancement during auditory descriptive naming. We anticipated that the distinct structures of auditory descriptive questions would help us identify which combinations of phrases drive functional connectivity enhancement within and between hemispheres. For example, comparing the neural dynamics when combining two concrete phrases versus a *wh*-interrogative and a concrete phrase is expected to reveal differences in language processing. The former more easily elicits a semantically relevant response, while the latter requires an additional stimulus phrase before a response can be formulated. Thus, we hypothesize that intra-hemispheric functional connectivity between the left temporal and frontal lobes intensify when patients combine two concrete phrases. We further anticipate that these functional connectivity enhancements will be at least in part supported by the arcuate fasciculus, which connects the temporal and frontal lobe neocortices (Catani et al., 2005; Hagoort and Indefrey, 2014; Gajardo-Vidal et al., 2021; Rocchi et al., 2021). A previous iEEG study reported that high-gamma amplitude in the left pIFG and MTG was augmented when patients were required to combine more words during a sentence reading task (Woolnough et al., 2023). Additionally, our previous iEEG study, which employed the same auditory descriptive naming task in 23 patients with focal epilepsy, found that the left prefrontal regions exhibited modest but significant high-gamma attenuation upon hearing a *wh-*interrogative at the onset of a question (Iwaki et al., 2021).

Furthermore, we predict that neural engagement in the left perisylvian neocortical regions, including the pIFG, will be more intense when patients are required to answer *what* questions compared to *when* or *where* questions. This prediction is based on the need for lexical selection from a broader range of potential responses to *what* questions. *When* is used to ask a time-related question, *where* is used to ask a place-related question, whereas *what* is used to ask questions related to time, place, and many other subjects. Previous lesion, fMRI, and iEEG studies have indicated that the left perisylvian neocortical regions, including the pIFG, are involved in lexical selection from a broad range of correct answers (Schnur et al., 2009; Cho-Hisamoto et al., 2015; Riès et al., 2015). Through our analyses, we expect to identify functional and effective connectivity pathways that are directly connected to the cortical regions exhibiting increased neural engagement in response to questions asking *what*. We anticipate that the resulting connectivity atlas will serve as a novel neurobiological model of auditory descriptive naming.

## 2. METHODS

### 2.1. Participants

The inclusion criteria for this study were native Japanese speakers with drug-resistant focal seizures who participated in an auditory descriptive naming task during iEEG recording sessions at either the National Center of Neurology and Psychiatry in Tokyo, Japan, or the Tohoku University Hospital in Sendai, Japan. The exclusion criteria included: (i) an inability to complete the task, (ii) significant brain malformations affecting the central or lateral sulcus, (iii) a history of prior resective epilepsy surgery, and (iv) evidence of right-hemispheric language dominance, as indicated by the Wada test results or left-handedness in conjunction with left-hemispheric congenital neocortical lesions. In other words, our eligibility criteria ensured that the left hemisphere of all patients contained essential language functions. In our previous iEEG study (Sonoda et al., 2022), we extensively discussed the validity of inferring right-hemispheric language dominance based on left-handedness in conjunction with left-hemispheric congenital neocortical lesions. The current study received approval from the ethical committees of the National Center of Neurology and Psychiatry and the Tohoku University Graduate School of Medicine, and informed consent was obtained from all adult participants or the guardians of pediatric participants. **Table S1** describes the detailed patient demographic information.

### 2.2. iEEG recording

Based on the results of non-invasive evaluations, we decided where to place subdural and depth platinum electrodes in the hemisphere impacted by epilepsy (with a spacing ranging from 5 to 10 mm between each). These evaluations included examining the seizure semiology, scalp video-EEG, and MRI. To estimate the epileptogenic zone for surgical removal, we continuously acquired iEEG signals at a rate of 1,000 Hz directly from the cerebral cortex using a Nihon Kohden EEG 1200 system (Nihon Kohden, Tokyo, Japan). To minimize the effect of pathological discharges on the measurement of event-related high-gamma activity (Mercier et al., 2022), we excluded electrode sites from further analysis if they were identified as part of the seizure onset zones (SOZs) triggering habitual seizures (Asano et al., 2009), irritative zones (those generating interictal epileptiform discharges; Kural et al., 2020), or areas impacted by MRI-visible structural lesions or artifacts.

### 2.3. Coregistration of intracranial electrodes and MRI

Using a preoperative T1-weighted spoiled gradient-echo volumetric MRI and post-electrode implant CT image, we generated a three-dimensional brain image showing the positions of the intracranial electrodes on the cerebral cortex (Stolk et al., 2018). Three board-certified neurosurgeons (K.S., H.U., and N.K.) verified the accuracy of the intracranial electrodes’ placement based on the visual assessment of intraoperative photographs, CT, and MRI (Kuroda et al., 2021). We used FreeSurfer scripts (http://surfer.nmr.mgh.harvard.edu) to standardize the electrode locations across individuals and projected these sites onto the average cortical surface image provided by FreeSurfer. Additionally, we parcellated the cortical gyri into anatomical ROIs (**Figure S1**) (Desikan et al., 2006; Ghosh et al., 2010; Sakakura et al., 2023), and this procedure allowed us to associate each electrode site with a specific ROI.

### 2.4. Auditory description naming task

Patients participated in the task (**Figure 1**) while undergoing iEEG recording at the bedside. We ensured that none of the patients had experienced a seizure in the two hours preceding the task. Each patient was asked to verbally respond to 96 three-phrase questions, which were articulated by a native Japanese-speaking male and transmitted through a speaker (Iwaki et al., 2021). The length of each question was set at 1.8 seconds. The sentences used in these questions were categorized into two types based on their structure: [a] those starting with a *wh*-interrogative and [b] those ending with a *wh*-interrogative. To control for the syllable count and occurrence frequency of three-phrase sentences within the two different structures, we used the National Institute for Japanese Language and Linguistics Database (https://pj.ninjal.ac.jp/corpus_center/bccwj/en/), ensuring there was no significant difference between them (two-sided p-value >0.05 on one-way ANOVA). We also counterbalanced the sentence structures among patients, leveraging the flexibility of Japanese language to express the same meaning in different phrase orders.

**Figure 1.**
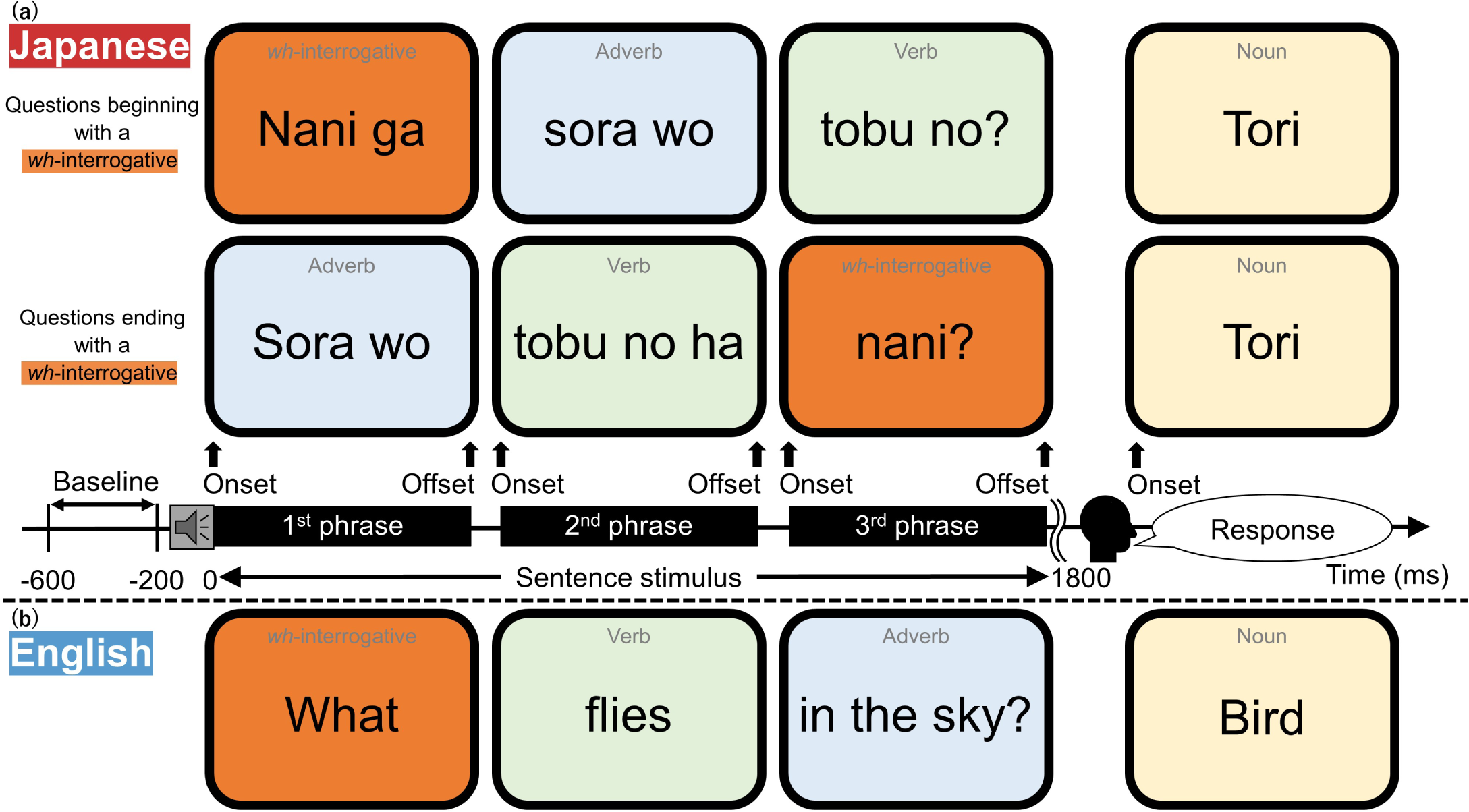
Auditory description naming task. (a) Here, we present that a question with the same meaning in Japanese (i.e., ‘What flies in the sky?’ in English) can be presented with two distinct structures, and both structures are used equally commonly by Japanese-speaking individuals. In the present study, each Japanese-speaking patient verbally answered each of the 96 three-phrase questions. Forty-eight questions started with a *wh-*interrogative followed by either an adverb or object, and then by a verb. The remaining 48 questions started with either an adverb or object, followed by a verb, and then by a *wh-*interrogative. (b) In English, this three-phrase question ‘What flies in the sky?’ can be given in this order: a *wh-*interrogative followed by a verb, and then by either an adverb or object.

[a] Questions starting with a *wh*-interrogative: In 48 trials, each question was structured to begin with a ‘*wh*-interrogative’, followed by an ‘adverb or object’, and then a ‘verb’. The *wh-* interrogative was either *what* (16 trials), *when* (16 trials), or *where* (16 trials). The durations were as follows: the first phrase averaged 436 ± 56 ms, the second phrase 487 ± 100 ms, and the third phrase 554 ± 75 ms (mean ± standard deviation). Regarding the intervals, there was an average gap of 181 ± 62 ms between the first and second phrases, and 157 ± 60 ms between the second and third phrases.

[b] Questions ending with a *wh*-interrogative: In the remaining 48 trials, in which questions ended with a *wh*-interrogative, each question was structured to begin with an ‘adverb or object’, followed by a ‘verb’, and then a ‘*wh*-interrogative’. In this structure, the initial adverb or object was a concrete noun, reflecting the Japanese sentence structure where an adposition follows a noun, as opposed to English where it precedes. The durations for these phrases were as follows: the first phrase averaged 546 ± 80 ms, the second phrase 538 ± 83 ms, and the third phrase 410 ± 50 ms (mean ± standard deviation). The intervals averaged 168 ± 57 ms between the first and second phrases, and 155 ± 49 ms between the second and third phrases.

The questions were presented in a pseudorandom order, ensuring that no two consecutive trials featured the same *wh*-interrogative. Response time was defined as the duration between the end of the question (i.e., the offset of the 3rd phrase) and the start of the patient’s response. Trials where a patient did not overtly verbalize a correct answer were excluded from the measurement of event-related high-gamma modulations.

### 2.5. Statistical assessment of the association between patient profiles and behaviors

Prior to surgery, we measured patients’ intelligence quotient (IQ) levels with the Wechsler Intelligence Scale. To determine the influence of patient demographics on patient’s behavioral responses, we utilized a mixed model analysis. The fixed effect factors in this model included patient age, full-scale IQ, the presence of a SOZ in the left hemisphere, and the number of antiseizure medications taken immediately prior to the iEEG recording, with response accuracy and median response time as the dependent variables. We interpret the use of multiple antiseizure medications as indicative of greater seizure-related cognitive impairment, as polytherapy often correlates with more severe seizures and cognitive burdens (Kwan and Brodie, 2001; Kuroda et al., 2021). The random effect factors included intercept and patient. We used SPSS Statistics 28 (IBM Corp., Chicago, IL, USA) for statistical analyses. All reported p-values are two-sided, and a p-value of <0.05 was considered statistically significant. A significant independent effect of the patient’s profile on response accuracy or response time, if present, would suggest that we need to assess its effect on high-gamma dynamics.

### 2.6. Statistical assessment of the influence of question types on behaviors

To assess the impact of different question structures on patient responses, we compared response accuracy and median response times across trials using the Wilcoxon Signed-Rank Test, focusing on differences between questions beginning and ending with a *wh*-interrogative. If the question structures affected response accuracy or response times, our assumption that sentence comprehension difficulty would be equivalent between the two distinct phrase orders would be deemed infeasible.

Additionally, within sentences sharing the same structure, we identified significant variations in response accuracy and response times to questions involving *what, when,* and *where,* using the Friedman test. If significant variations were found, a pairwise comparison was performed using the Wilcoxon Signed-Rank Test. A p-value of <0.05 was considered statistically significant. If *what* questions showed lower response accuracy or longer response times compared to *when* or *where* questions, our notion that *what* questions require more neural resources for lexical selection from a broader range of potential responses would be supported.

### 2.7. Measurement of cortical high-gamma modulations

We measured task-related high-gamma modulations using a time-frequency analysis reported in our previous studies (Nakai et al., 2017; Kitazawa et al., 2023). The process involved transforming iEEG signals to the time-frequency domain, with increments of 10 ms and 5 Hz, through a complex demodulation technique (Papp and Ktonas, 1977; Hoechstetter et al., 2004). This technique, part of the BESA EEG software package (BESA GmbH, Gräfelfing, Germany), involved first multiplying the time-domain iEEG signal with a complex exponential, then applying a band-pass filter. The complex demodulation, which uses a Gaussian-shaped low-pass finite impulse response filter, essentially performs a Gabor transformation. We measured high-gamma_70-110 Hz_ amplitude changes as a percent change relative to the resting baseline (-600 to-200 ms pre-question onset), at each 10-ms/5-Hz time-frequency bin. The time-frequency resolution for these high-gamma measurements was approximately ±15.8 ms and ±7.1 Hz, defined by the 50% power reduction point of the filter. We performed this time-frequency analysis using a bipolar montage technique to minimize contamination from volume conduction by measuring the potential difference between adjacent electrode pairs. For each cortical electrode site that is flanked by two or more neighboring pairs, we calculated the high-gamma amplitude value by averaging the measurements from these pairs. For instance, the high-gamma amplitude value for the cortical electrode site A2 on a depth electrode was the average between those from the A1-A2 and A2-A3 pairs (Johnson et al., 2023). This measurement was time-locked to the onset and offset of each stimulus phrase and to the onset of the patient’s response. We created an animated movie to illustrate the percent change in high-gamma amplitudes, measured within 10 mm from the center of each electrode and averaged across electrodes from all patients. We then displayed these dynamics of high-gamma activity on the FreeSurfer standard brain template.

### 2.8. Statistical assessment of cortical high-gamma modulations

We determined the time bins and ROIs where significant task-related high-gamma modulations occurred, using the method reported in our previous studies (Nakai et al., 2019; Kitazawa et al., 2023). In each ROI, we plotted the mean task-related high-gamma amplitude, accompanied by a 99.99% confidence interval (99.99%CI) shading. This was done for all cortical electrode sites that were free from artifacts and outside the SOZs, irritative zones and MRI structural lesions. This analysis included all ROIs that had at least five cortical electrode sites derived from at least three patients in each hemisphere. **Table S2** lists the numbers of cortical electrode sites and patients within each of the 40 ROIs (20 in each hemisphere) that met this requirement and were included in this analysis. We used a one-sample t-test to identify the moments and specific ROIs where task-related high-gamma amplitudes significantly deviated from the baseline. A significant augmentation of task-related high-gamma activity was determined when the lower limit of the 99.99%CI for the mean amplitude consistently exceeded zero for a duration of 50 ms or more. Similarly, a significant attenuation was noted when the upper limit of the 99.99%CI consistently fell below zero for at least 50 ms. Using a 99.99%CI corresponds to a Bonferroni adjustment for 500 repeated tests, whereas a 50-ms interval would include a minimum of three high-gamma oscillatory cycles. While this stringent significance criterion raises the likelihood of a Type II error, it simultaneously decreases the possibility of a Type I error. Below, we used the timing of significant high-gamma augmentation and attenuation at ROIs to compute the dynamics of functional connectivity modulations.

### 2.9. Statistical assessment of the influence of patient profiles on cortical high-gamma modulations

To evaluate the influence of patient profiles on task-related cortical high-gamma modulations, we conducted a mixed model analysis focusing on each specific 300-ms epoch between the question onset and the patient’s response. This analysis incorporated fixed effect factors including patient age, full-scale IQ, the presence of a SOZ in the left hemisphere, and the number of antiseizure medications administered immediately before the iEEG recording. The dependent variable in this model was the amplitude of high-gamma activity at given ROIs. Given the examination of 13 distinct 300-ms epochs (i.e., before and after phrase onsets and offset as well as response onset) at 40 ROIs for high-gamma amplitude measurements within this framework, we applied a Bonferroni correction to adjust for multiple comparisons, considering a corrected p-value of <0.05 as statistically significant.

### 2.10. Statistical assessment of the influence of question types on cortical high-gamma modulations

We likewise identified the time bins and ROIs where high-gamma amplitudes significantly differed between questions beginning with *what, when,* and *where*, using a one-sample t-test. We defined significant differences in high-gamma amplitudes at given ROIs when the lower limit of the 99.99%CI of the mean difference in high-gamma amplitude was greater than zero for at least 50 ms consecutively. Similarly, we analyzed the timing and locations of high-gamma amplitude variations in questions ending with *what, when,* and *where.* We predicted that high-gamma amplitudes in the left perisylvian neocortical regions would be greater for *what* questions compared to *when* or *where* questions. Such preferential high-gamma augmentation for *what* questions, if occurring between the question offset and response, cannot be simply attributed to differences in syllable count or occurrence frequency of stimuli across question types. Therefore, the analysis below also aimed to examine the characteristics of functional and effective connectivity modulations associated with an ROI showing this specific pattern of high-gamma augmentation in response to *what* questions.

### 2.11. Measurement of functional connectivity modulations

We determined the time bins and white matter pathways where functional connectivity was significantly modulated using a method similar to those reported in our previous studies (Kitazawa et al., 2023; Ono et al., 2023; Ueda et al., 2024). We used diffusion-weighted imaging (DWI) tractography data to visualize the dynamic functional connectivity in white matter between cortical ROIs. We defined functional connectivity between cortical ROIs as either significantly enhanced or diminished based on two criteria: (1) both cortical ROIs must exhibit significant, simultaneous, and sustained changes in high-gamma amplitude for at least 50 ms, and (2) these regions need to be connected by direct DWI streamlines. This duration corresponds to five consecutive 10-ms time bins. To assess the likelihood of random co-occurrence of high-gamma augmentation (i.e., Type I error), we calculated the chance probability of co-augmentation lasting at least 50 ms when significant high-gamma augmentation occurred in χ% of the entire analysis period on average across 40 cortical ROIs. Given the 780 distinct pairs of analysis ROIs in the present study, the chance probability of simultaneous occurrence of significant high-gamma co-augmentation at least at a pair of ROIs for consecutive five 10-ms time windows was estimated based on the following equation: 1 – (1 – ((χ/100)^2^)^5^)^780^ (Ueda et al., 2024).

### 2.12. Visualization of functional connectivity through white matter

Neuroscientists often rely on brain templates provided by the Montreal Neurological Institute (MNI) or FreeSurfer, working under the assumption that these templates reflect the brain of an average patient with a reasonable accuracy. In the present study, we adopted a similar stance, suggesting that the white matter streamlines in both patients and healthy individuals are broadly similar. Using open-source DWI data from 1,065 healthy individuals (Yeh et al., 2018; http://brain.labsolver.org/diffusion-mri-templates/hcp-842-hcp-1021), we mapped white matter streamlines, replicating methods previously employed (Kitazawa et al., 2023; Sakakura et al., 2023). The validity of employing the open-source DWI alongside individual iEEG data for exploring human brain network dynamics has been confirmed by two independent groups (Mitsuhashi et al., 2021; Sonoda et al., 2021; Azeem et al., 2023). Seeds were placed in cortical ROIs that showed significant high-gamma co-augmentation (or co-attenuation) for at least 50 ms. We utilized the DSI Studio script (http://dsi-studio.labsolver.org/) to visualize tractography streamlines that directly connect these ROIs within the MNI standard space. The parameters set for fiber tracking included a quantitative anisotropy threshold of 0.05, a maximum turning angle of 70°, and a streamline length ranging from 20 to 250 mm. Streamlines involving the brainstem, basal ganglia, thalamus, or cerebrospinal fluid space were not included in our dynamic tractography analysis. Our resulting dynamic tractography video atlases illustrate functional connectivity modulations, captured in 10-ms increments across each 50-ms epoch. We also provided the dynamic connectome matrices that visualize the spatiotemporal characteristics of cortical ROI-pairs with significant functional connectivity enhancement and diminution during each time period.

### 2.13. Statistical assessment of the influence of question structures on functional connectivity

We assessed the impact of question structures on the spatial extent of functional connectivity by comparing the proportion of ROI pairs with significant functional connectivity enhancement among all ROI pairs. To this end, we employed a 2×2 Fisher’s exact test, focusing specifically on questions that either begin or end with a *wh-*interrogative. This analysis was conducted at each 10-ms time bin separately for intra-hemispheric pathways within each hemisphere and for inter-hemispheric pathways between hemispheres. A significant difference in proportions, indicated by a p-value of <0.05 across five consecutive 10-ms time windows, would suggest that the question structure significantly affects the spatial extent of functional connectivity. Our hypothesis posits that functional connectivity within the left perisylvian networks is enhanced when humans are required to combine two concrete phrases compared to when combining a *wh-*interrogative with a concrete phrase and that the left arcuate fasciculus contributes to such functional connectivity enhancement. Consequently, we predicted that, upon hearing the second phrase, the proportion of significant intra-hemispheric functional connectivity enhancement within the left hemisphere would be greater for questions ending with a *wh-*interrogative (i.e., those beginning with’adverb or object’ followed by’verb’) than for those beginning with a *wh-*interrogative followed by’adverb or object’.

### 2.14. Measurement of effective connectivity

We determined the white matter pathways and direction of neural information flows using a transfer entropy-based effective connectivity analysis, similar to that previously reported (Firestone et al., 2023). We conducted analyses at given 200-ms epochs (sliding every 100 ms and each including twenty 10-ms time bins) between the first phrase onset and overt response. We declared facilitatory information flow from ROI_1_ to ROI_2_ during a given 200-ms epoch when the following three criteria were met: [1] ROI_1_ and ROI_2_ were connected by a direct white matter tract on DWI tractography. [2] Functional connectivity between ROI_1_ and ROI_2_ was significantly enhanced through the white matter tract during the epoch of interest (as detailed in the preceding analysis). [3] Transfer entropy indicated the presence of a neural information flow (i.e., above zero) between ROIs showing significantly enhanced functional connectivity. Transfer entropy was calculated using a custom Matlab script adapted from the algorithm described in Ito et al., 2011. Time-frequency data was binarized for input into the transfer entropy program. Thereby, each 10-ms time bin was assigned a value of 1 if the high-gamma amplitude across all available electrode sites within a given ROI was significantly higher (p-value <0.05) than the mean during the immediately preceding 100 ms, as determined by a paired t-test; otherwise, it was assigned a value of 0. Since transfer entropy inherently measures conditional mutual information (Schreiber 2000) in’bits’, it was necessary to binarize the continuous high-gamma amplitude data to make it compatible with the equation. The strength of information flows, both efferent (arising from the ROI) and afferent (converging to the ROI), was computed at each ROI.

We likewise computed the strength of suppressive neural information flows between given ROIs through white matter pathways showing functional connectivity diminution. To this end, we used a binary time series of high-gamma amplitude data in which each 10-ms time bin was assigned a value of 1 if the high-gamma amplitude within a given ROI was significantly lower than the mean during the immediately preceding 100 ms; otherwise, it was assigned a value of 0.

### 2.15. Statistical assessment on effective connectivity

We determined during which 200-ms epochs the strengths of neural information flows across the left hemispheric ROI pairs became higher compared to those within the right hemisphere, using the Wilcoxon Signed-Rank Test. We predicted that the left intra-hemispheric information flows would be higher than those on the right around the third phrase offset and before response onset, when participants are anticipated to generate a verbal response internally. Subsequently, we determined during which 200-ms epochs the strength of information flows across the left hemispheric ROI pairs became different between questions beginning and ending with a *wh-* interrogative. We predicted that the left intra-hemispheric flows would be higher when participants were required to combine two concrete phrases, more so than when combining a wh-interrogative and a concrete phrase. Since information flows were measured at 41 200-ms epochs, we treated a Bonferroni-corrected p-value of <0.05 as significant.

All of our connectivity analyses, as described above, focused on assessing relationships between pairs of specific ROIs; interactions among multiple ROIs were not analyzed simultaneously.

### 2.16. Data availability

The iEEG data are available at https://openneuro.org/datasets/ds005007/versions/1.0.0 (doi:10.18112/openneuro.ds005007.v1.0.0).

### 2.17. Code availability

The analysis codes are available at https://github.com/md462588/Demo.git (doi: 10.5281/zenodo.10828427).

## 3. RESULTS

### 3.1. Patient profiles

A total of 40 Japanese patients (16 females) satisfied the eligibility criteria, completed the auditory description naming task, and were included in the study (**Table S1**). The mean age was 25 years (range: 6-54). The mean full-scale IQ was 79.6 (range: 50-114). The SOZ involved the left hemisphere in 22 patients. The mean number of oral antiseizure medications taken immediately prior to the iEEG recording was 3.1 (range: 1-5). The total number of artifact-free, non-epileptic electrode sites was 747 (median: 27 per patient; interquartile range [IQR]: 9 to 47) in the left hemisphere and 822 (median: 28 per patient; IQR: 9 to 51) in the right hemisphere.

### 3.2. Behavioral observations

The median correct response rate was 0.94 across 40 patients (range: 0.72 to 1.00). The mixed model analysis demonstrated that older patient age was associated with higher correct response rates (mixed model estimate: 0.003; degree of freedom [DF]: 34.994; t-value: 2.250; p-value: 0.031; 95%CI: 0.0003 to 0.005). None of the other fixed effect variables showed significant association with the response rate (p-values: > 0.134). The median response time was 1,574 ms across 40 patients (standard deviation: 1,033 ms). The mixed model analysis did not find an association between the response time and any of the fixed effect variables (p-values: >0.424).

The Wilcoxon Signed-Rank Test did not find a significant difference in correct response rate (p-value: 0.13) or response time (p-value: 0.71) between questions beginning and ending with a *wh*-interrogative. The Friedman test found a significant difference in correct response rate (p-value: 0.007) but not response time (p-value: 0.38) across questions asking *what, when,* and *where*. The median correct response rates for *what, when,* and *where* questions were 92.2%, 96.9%, and 93.8%, respectively. A post-hoc Wilcoxon Signed-Rank Test revealed that the correct response rate for *what* questions was significantly lower compared to *when* (p-value: <0.001) and *where* questions (p-value: 0.014).

### 3.3. Summary description of the functional and effective connectivity atlas of auditory naming

Prior to presenting specific statistical results, we provide a visual overview of the spatiotemporal dynamics observed in high-gamma amplitude modulations, functional connectivity, and effective connectivity during an auditory description naming task involving overt responses to three-phrase questions consisting of a *wh-*interrogative followed by either an adverb or object, then followed by a verb. Detailed visual representations are provided in **Video S1**, while **Video S2** illustrates the temporal changes in anatomical ROI pairs exhibiting functional connectivity modulations and directional neural information flows.

Upon hearing a *wh-*interrogative as the onset of questions, we observed high-gamma augmentation in the bilateral STG, enhanced inter-hemispheric functional connectivity between the left and right STG via the posterior corpus callosum, and facilitatory neural information flows from the left to right STG (**Figure 2A**). Conversely, inter-hemispheric functional connectivity decreased between the right and left STG through the anterior corpus callosum, accompanied by suppressive neural information flows from the right to left anterior middle frontal gyrus (aMFG) (**Figure 2B**). Right-hemispheric inter-hemispheric functional connectivity also decreased between the aMFG and superior parietal gyrus (SPG) through the superior longitudinal fasciculus, accompanied by bidirectional, suppressive neural information flows between them.

**Figure 2.**
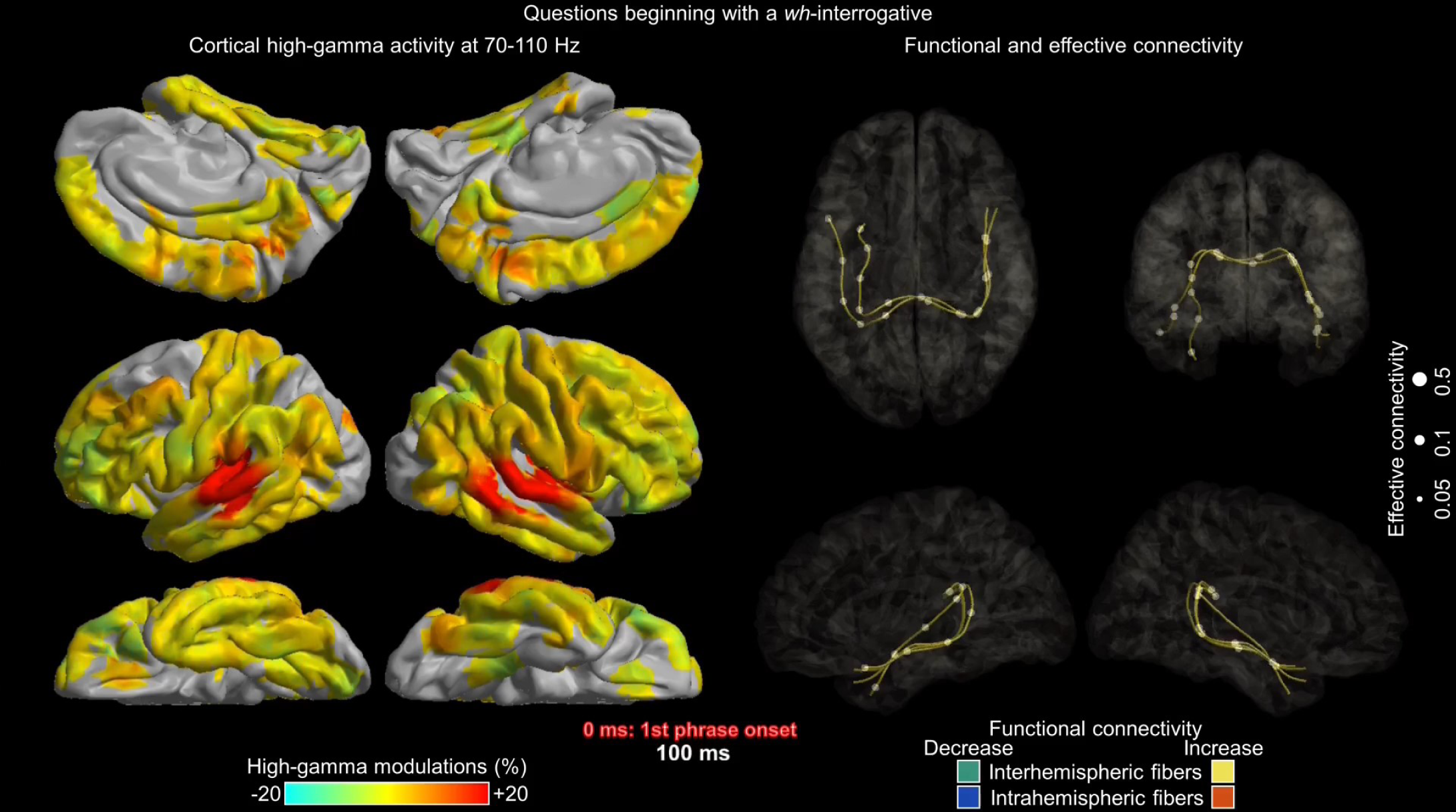
Functional and effective connectivity atlas of naming responses to questions beginning with a *wh-*interrogative. The snapshots on the left demonstrate the dynamics of cortical high-gamma amplitude modulations, while those on the right illustrate the dynamics of functional and effective connectivity during an auditory description naming task involving overt responses to three-phrase questions beginning with a *wh-*interrogative, followed by either an adverb or object, and then a verb. Yellow white matter streamlines indicate significant interhemispheric functional connectivity enhancement, with orange indicating significant intrahemispheric enhancement. Conversely, green streamlines show significant inter-hemispheric connectivity diminution, and blue ones indicate intra-hemispheric diminution. The strength of neural information flows is represented by the size of the white circles. The direction of neural information flows are visually indicated by moving circles on **Video S1** and triangle locations on **Video S2**. (A) 100 ms after the first phrase onset. (B) 400 ms after the first phrase onset. (C) 300 ms prior to the third phrase offset. (D) 350 ms prior to the response onset. (E) 100 ms after the response onset.

While hearing the second phrase consisting of either an adverb or object and the third phrase consisting of a verb, high-gamma augmentation persisted in the bilateral STG. Inter-hemispheric functional connectivity remained enhanced between the left and right STG, with facilitatory neural information flows observed in both directions.

At the offset of the third phrase (i.e., at question offset), high-gamma augmentation became prominent in the extensive perisylvian cortical regions within the left hemisphere. Left intra-hemispheric functional connectivity enhancement mainly through the arcuate fasciculus became more prominent than inter-hemispheric functional connectivity enhancement, and bidirectional, facilitatory neural information flows took place between the left temporal and frontal regions (**Figure 2C**).

As the response onset approached, high-gamma augmentation became prominent in the left prefrontal, premotor, and precentral regions. Intra-hemispheric functional connectivity was heightened across these left frontal lobe regions, and bidirectional, facilitatory neural information flows were observed within these regions (**Figure 2D**).

Immediately prior to and during responses, high-gamma augmentation was observed in the bilateral precentral, postcentral, and STG regions. Inter-hemispheric functional connectivity was enhanced across precentral and postcentral gyri, as well as between bilateral STG through the posterior corpus callosum, with bidirectional, facilitatory neural information flows occurring between these regions (**Figure 2E**).

**Videos S3 and S4** likewise provide detailed visual representations of the functional and effective connectivity concerning naming responses to three-phrase questions with a different phrase order. These questions begin with either an adverb or object, followed by a verb, and conclude with a *wh-*interrogative. Overall, the high-gamma activity and connectivity patterns exhibited similarities between questions that began and ended with a *wh-*interrogative. However, notable differences were observed, particularly the absence of inter-hemispheric functional connectivity diminution or suppressive neural information flows between the right and left prefrontal regions (see 00:14-00:17 on **Video S4**). Instead, an earlier emergence of enhanced intra-hemispheric functional connectivity and bidirectional neural information flows through the left arcuate fasciculus were observed while hearing the second phrase (see 00:37 on **Video S4**).

### 3.4. Statistical assessment of the effects of patient profiles on cortical high-gamma modulations

Individuals with higher IQ scores exhibited less lingering high-gamma augmentation in the left posterior MFG (pMFG) during responses. The mixed model analysis revealed that a higher full-scale IQ was independently associated with decreased high-gamma amplitudes in the left pMFG during the 300-ms period immediately following the response onset (mixed model estimate:-1.1%; DF: 29; t-value:-6.27; uncorrected p-value: 0.0000008; 95%CI:-1.4% to-0.7%). We failed to find that age, the presence of a SOZ in the left hemisphere, or the number of antiseizure medications significantly affected high-gamma amplitude at any ROIs, with all Bonferroni corrected p-values remaining above 0.05.

### 3.5. Statistical comparisons of intra-hemispheric functional connectivity between the left and right hemispheres

**Video S5** illustrates the dynamics of cortical high-gamma amplitude at each ROI during naming responses to questions beginning and ending with a *wh-*interrogative. **Video S6** shows the dynamic changes in the spatial extent of (i.e., the number of ROI pairs showing) functional connectivity enhancement and diminution. The Type I error rate for detecting functional connectivity enhancement between at least one pair of ROIs was estimated to be very small, at most 0.00000006, in this study. More precisely, significant high-gamma augmentation was observed in 9.8% of the entire analysis periods, on average, across 40 cortical ROIs. The chance probability of randomly observing co-augmentation of high-gamma amplitudes lasting ≥50 ms between at least one pair of ROIs was approximately 0.00000006.

In questions starting and ending with a *wh*-interrogative, we observed a notable pattern: the spatial extent of intra-hemispheric functional connectivity enhancement was significantly greater in the left hemisphere than in the right during specific periods. This left-hemispheric dominant pattern was prominent both [1] upon hearing a verb following either an adverb or an object and [2] prior to the patient’s response (**Figure 3A**).

**Figure 3:**
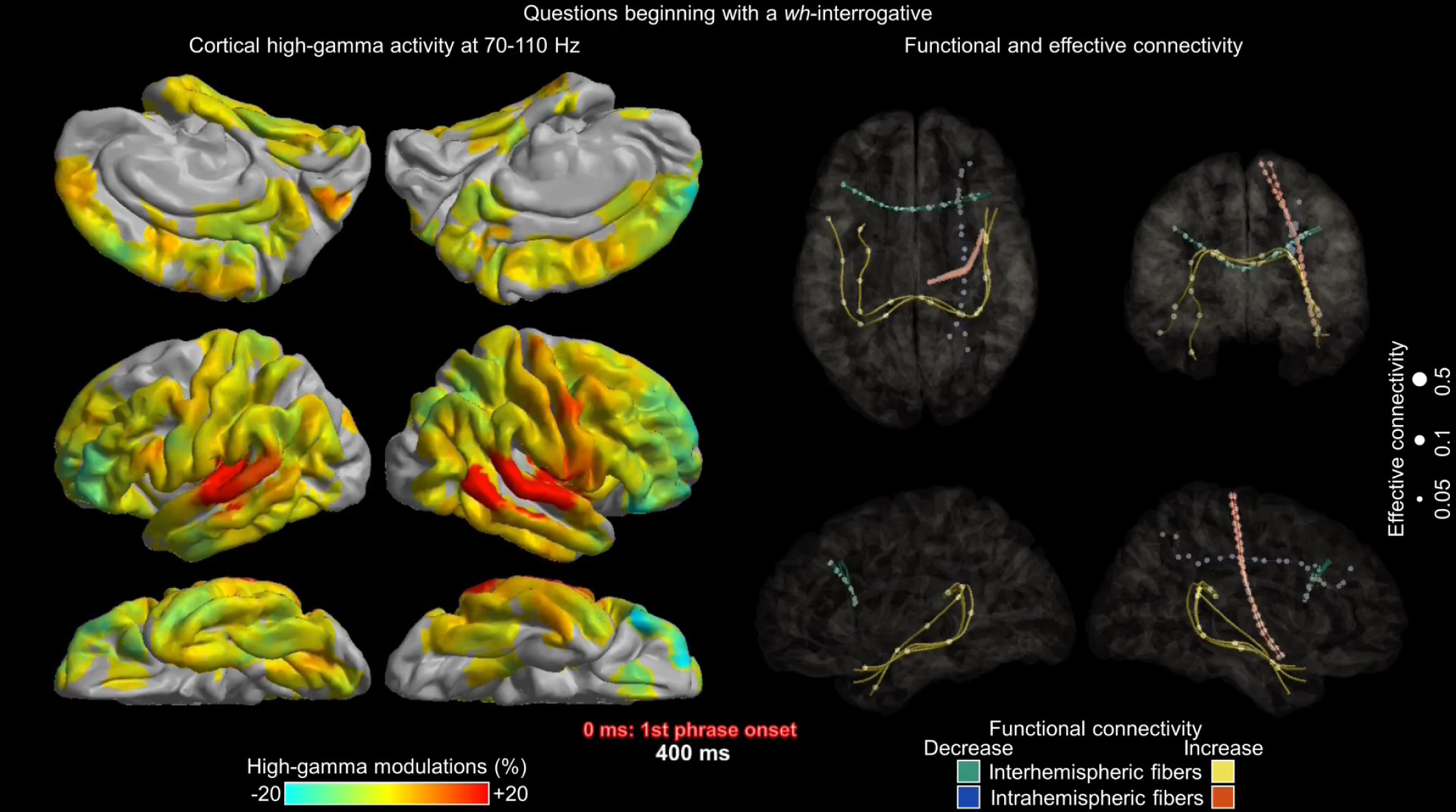
Functional connectivity dynamics during naming responses. (A) This figure demonstrates the dynamic changes in the spatial extent of intra-hemispheric functional connectivity enhancement, represented by the number of region-of-interest (ROI) pairs, during the process of naming responses. The orange line represents the left hemisphere and purple represents the right hemisphere. A horizontal bar marks the time points at which the proportion of ROI pairs showing significant functional connectivity enhancement was significantly higher in the left hemisphere than in the right. (B) This figure demonstrates the dynamic changes in the spatial extent of functional connectivity enhancement during the process of responding to questions beginning (green lines) and ending (purple lines) with a *wh*-interrogative. A horizontal bar marks the time points at which the proportion of ROI pairs showing significant functional connectivity enhancement was significantly higher in questions ending rather than beginning with a *wh*interrogative.

More precisely, for questions starting with a *wh-*interrogative, the proportion of ROI pairs showing significant functional connectivity enhancement within the left hemisphere exceeded that in the right hemisphere during the-350 to-210 ms <u>pre-third phrase offset</u> (i.e., pre-verb offset) (Fisher exact probability test p-values: ≤0.03 for a consecutive 140-ms period; maximum effect size h: 0.463). During this period, up to ten left intra-hemispheric white matter pathways exhibited functional connectivity enhancement, in contrast to essentially zero in the right hemisphere. Similarly, during 310-ms periods between-400 and +30 ms relative to the response onset, the left hemisphere demonstrated a greater proportion of ROI pairs with significant functional connectivity enhancement compared to the right hemisphere (Fisher exact probability test p-values: ≤0.02 for ≥50-ms periods; maximum effect size h: 0.480). During these periods, up to 18 left intra-hemispheric pathways exhibited functional connectivity enhancement, in contrast to at most three in the right hemisphere (**Figure 3A**).

For questions ending with a *wh-*interrogative, the proportion of ROI pairs showing significant functional connectivity enhancement within the left hemisphere exceeded that in the right hemisphere during the-100 to-50 ms <u>pre-second phrase offset</u> (i.e., pre-verb offset) (Fisher exact probability test p-values: 0.03 for a consecutive 50-ms period; maximum effect size h: 0.357). During this period, six left intra-hemispheric white matter pathways exhibited functional connectivity enhancement, in contrast to none in the right hemisphere. Similarly, during 280-ms periods between-390 and-20 ms relative to the response onset, the left hemisphere demonstrated a greater proportion of ROI pairs with significant functional connectivity enhancement compared to the right hemisphere (Fisher exact probability test p-values: ≤0.03 for ≥50-ms periods; maximum effect size h: 0.409). During these periods, up to 20 left intra-hemispheric pathways exhibited functional connectivity enhancement, in contrast to at most three in the right hemisphere (**Figure 3A**).

The aforementioned left-hemispheric white matter pathways showing functional connectivity enhancement included the arcuate fasciculus connecting the left temporal and frontal lobe regions, the superior longitudinal fasciculus connecting the left frontal and parietal lobe regions, and the frontal lobe u-fibers and frontal aslant fasciculus connecting the left prefrontal, premotor, and Rolandic regions (see 00:12-00:20 on **Video S1**).

### 3.6. Statistical comparisons of intra-hemispheric effective connectivity between the left and right hemispheres

**Videos S2 and S4** present the directions of facilitatory and suppressive information flows between ROIs: **Video S2** is dedicated to naming responses to questions beginning with a *wh-*interrogative, while **Video S4** examines naming responses to questions ending with a *wh-*interrogative.

In questions starting with a *wh*-interrogative, we observed that the strength of the neural information flows was significantly greater in the left hemisphere than in the right during a 200-ms period-200 to-400 ms pre-third phrase offset (i.e., verb offset) (maximum neural information flow: 0.345 in the left and 0.032 in the right hemisphere; uncorrected p-value: 0.00016 on Wilcoxon Signed Rank Test). Neural information flows during this period were found between 10 ROI pairs within the left hemisphere, and all flows were bidirectional (see 01:12 on **Video S2**). The maximum neural information flow during this period was transmitted through the left arcuate fasciculus connecting the pMFG and ITG.

In questions ending with a *wh*-interrogative, our statistical analysis failed to demonstrate that the strength of neural information flows significantly differed between the left and right hemispheres. The maximum difference in the strength of neural information flows between the left and right hemispheres was noted during a 200-ms period 300-500 ms after the third phrase onset (i.e., *wh-* interrogative onset) (maximum neural information flow: 0.345 in the left and 0 in the right hemisphere; uncorrected p-value: 0.0039 on Wilcoxon Signed Rank Test) (see 01:04 on **Video S4**). this uncorrected p-value did not survive the Bonferroni correction.

### 3.7. Difference in neural dynamics between questions beginning and ending with a *wh*-interrogative

Our statistical analysis revealed that the spatial extent of left intra-hemispheric functional connectivity enhancement was larger when patients heard a concrete phrase following another concrete one, compared to hearing a concrete phrase following a *wh-*interrogative. More precisely, during a 50-ms period preceding the offset of the second phrase (i.e., prior to the third phrase), the proportion of ROI pairs exhibiting significant functional connectivity enhancement among all left-hemispheric ROI pairs was higher in questions beginning with two concrete phrases and ending with a *wh-*interrogative, compared to those beginning with a *wh-*interrogative followed by two concrete phrases (Fisher exact probability test p-values: 0.03; for a consecutive 50-ms period; maximum effect size h: 0.357; **Figure 3B**). In questions ending with a *wh-*interrogative, six out of the 190 left-hemispheric ROI pairs demonstrated functional connectivity enhancement, during this 50-ms period (i.e., upon hearing a verb following an adverb or object) (**Figure 3B**). The underlying white matter pathways included the left arcuate fasciculus, which connects the left STG with the left pIFG, the pMFG, and the precentral gyrus. Among these white matter pathways showing functional connectivity enhancement during this period, we noted bidirectional, facilitatory neural information flows between the left pIFG and precentral gyrus (see 00:50 on **Video S4**). Conversely, in questions beginning with a *wh-*interrogative, none of the left-hemispheric ROI pairs showed functional connectivity enhancement upon hearing the second phrase (i.e., upon hearing an adverb or object after an a *wh*-interrogative) (**Figure 3B**).

### 3.8. Statistical comparisons of neural dynamics across naming responses to questions asking *what*, *when*, and *where*

**Video S7** illustrates the differences in high-gamma amplitude between questions asking *what* and *when*, as well as between those asking *what* and *where*. We found that, in questions beginning with a *wh*-interrogative, high-gamma amplitude in the left pIFG around the offset of the third phrase (i.e., verb) was higher in questions asking *what* compared to in those asking *where* (**Figure 4A**). The maximum difference in high-gamma amplitude was observed to be 10.5%, with a 99.99%CI of 3.2% to 17.8% at-120 ms pre-third phrase offset. During a 200-ms period between-100 and - 300 ms pre-third phrase offset, the left pIFG showed enhanced functional connectivity with the left MTG and STG through the arcuate fasciculus, as well as with the left pMFG through the frontal lobe u-fibers (**Figure 4B-4C**). During the same 200-ms period, bidirectional, neural information flows took place across the left pIFG, MTG, and pMFG **Figure 4B-4C**).

**Figure 4:**
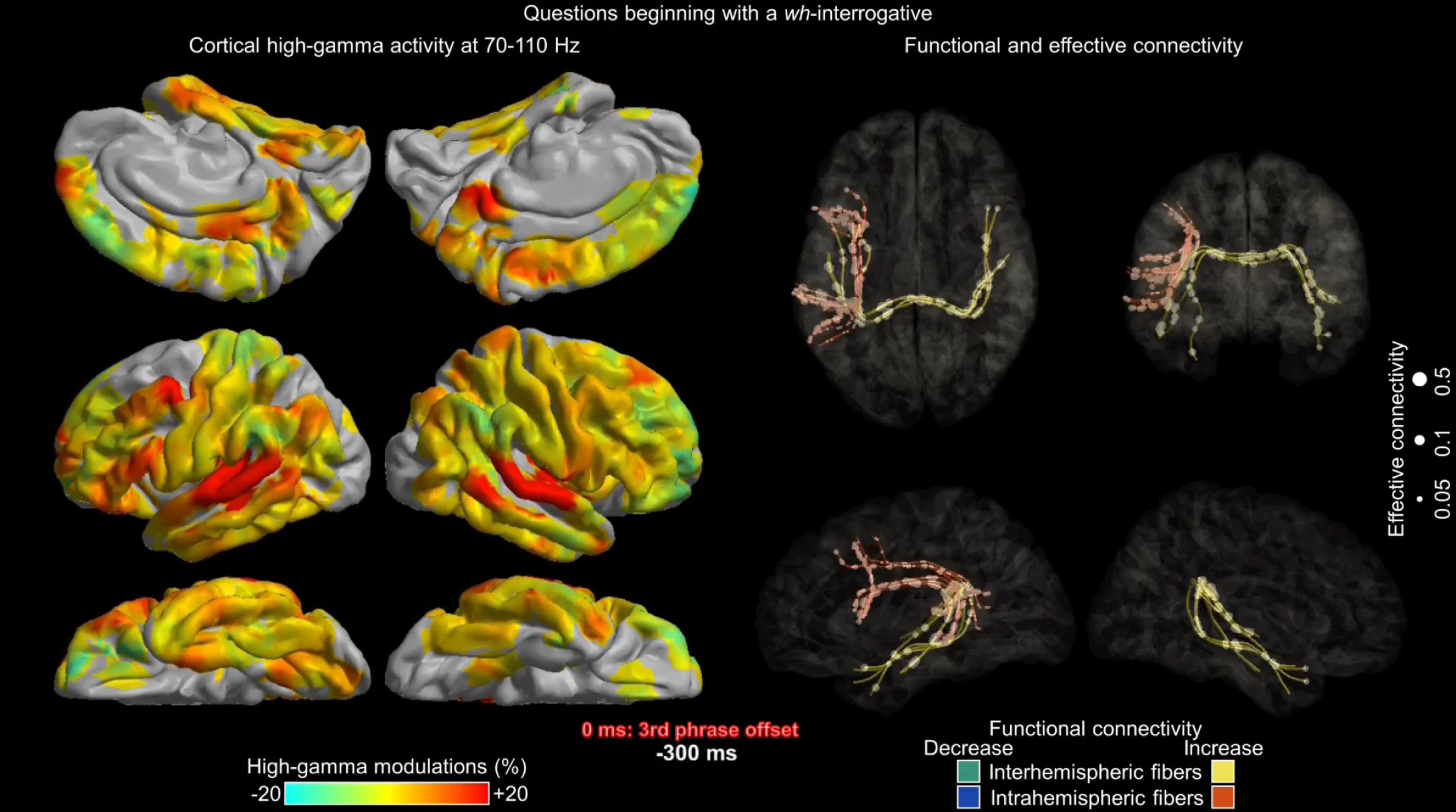
**Neural dynamics during naming responses to questions asking *what, when, and where.*** (A) This figure illustrates the dynamics of cortical high-gamma modulations in the left posterior inferior frontal gyrus (pIFG) in response to questions that begin with a *wh-*interrogative. It shows the mean high-gamma amplitudes along with their standard error shading. Blue line: questions beginning with *what*. Green lines: those beginning with *when*. Magenta line: those beginning with *where*. A horizontal bar indicates a time period where high-gamma amplitudes significantly differed between question types. (B) This matrix details the spatial extent of functional connectivity enhancement and neural information flows in trials in which patients were asked questions starting with *what*. During a 200-ms period-100 to-300 ms pre-third stimulus offset (i.e., verb), the left pIFG demonstrates enhanced functional connectivity with the left middle temporal, superior temporal, and posterior middle frontal gyri. (C) This figure displays the spatial characteristics of cortical high-gamma amplitudes, functional connectivity enhancement, and neural information flows in trials involving questions beginning with *what*. Specifically, it outlines the white matter pathways showing significant functional connectivity enhancement and neural information flows occurring within the same 200-ms period.

### 3.9. Robustness of interhemispheric facilitatory information flows

Using Spearman’s correlation tests, we confirmed that group-level high-gamma dynamics computed after excluding individual patients were consistent with results derived from the full dataset (**Table S3**). Additionally, we verified that interhemispheric facilitatory neural information flows connecting homotopic regions remained replicable after excluding either the patient with the shortest or longest response time (**Table S4**). These analyses demonstrate that the observed interhemispheric neural information flows preceding responses are not confounded by inter-patient variability in response times.

## 4. DISCUSSION

### 4.1. Connectivity modulations upon hearing a *wh*-interrogative at question onset

The present study aimed to visualize the dynamics of naming-related functional connectivity modulations on the order of hundreds of milliseconds, along with the strength and direction of neural information flows through direct white matter pathways. Such network measures were not assessed in our previous iEEG study (Iwaki et al., 2021), which focused solely on measuring local task-related high-gamma modulations in a given ROI. The resulting movies from the current study are novel and propose a neurobiological framework that may enhance our understanding of auditory naming processing in the human brain.

While participants were listening to a *wh-*interrogative (*what, when,* or *where*) marking the beginning of three-phrase descriptive questions, the bilateral STG engaged in a sustained manner (see 00:07-00:09 on **Video S1**). Such STG activation is expected to be involved in coordinated processing within the STG through local, short-range connections to interpret acoustic stimuli as speech sounds (Chang et al., 2010; Trumpis et al., 2021; Bhaya-Grossman and Chang, 2022; Norman-Haignere et al., 2022; Leonard et al., 2024). This sustained bilateral STG activation featured an enhancement of inter-hemispheric functional connectivity between these gyri via the posterior corpus callosum, and this was accompanied by facilitatory neural information flows initially from the left to the right STG and subsequently bidirectional (see 00:07-00:14 on **Video S1** and 00:11-01:12 on **Video S2**). The aforementioned observations are consistent with those from our prior iEEG study of 13 English-speaking patients, which also showed enhanced functional connectivity between the STG and the posterior corpus callosum during the listening of a *wh*-question (Kitazawa et al., 2023). Our observation of the left-to-right neural information flows can be in part attributed to the earlier onset (left: 40 ms vs right: 50 ms after the first phrase onset) and peak latency (left: 200 ms vs right 230 ms) of high-gamma amplitude augmentation in the left STG as compared to in the right STG (see 00:09 on **Video S5**). Our observations are in line with the notion that the left STG critically plays a role in analyzing temporal speech features, including the timing and rhythm of speech sounds essential for decoding the basic structure of speech, such as identifying phonemes and syllables and that temporal analysis is a prerequisite for prosody analysis in the hierarchical processing of verbal communication (Zatorre and Belin, 2001; Chang et al., 2010; Floegel et al., 2020). Conversely, the right STG is suggested to play a role in analyzing spectral features, including the tone and pitch fundamental to interpreting prosody (Zatorre and Belin, 2001; Floegel et al., 2020). It is plausible to hypothesize that the right STG exerts such spectral analysis in rapid succession of mental representations of speech sounds delivered from the left STG. A prior study using fMRI and DWI tractography provided observations consistent with our hypothesis (Elmer et al., 2016). Specifically, the study involved 13 professional musicians and 13 control participants and reported that the left and right auditory cortices are directly connected through the posterior corpus callosum. Furthermore, anatomical connectivity between the auditory cortices was positively correlated with performance in a phoneme categorization task, the degree of musical aptitude, and the intensity of hemodynamic responses in the left auditory cortex. The critical role of the posterior corpus callosum in auditory naming was suggested by a study of patients with drug-resistant epilepsy who underwent callosotomy and showed that postoperative speech impairment was severe when disconnection involved the posterior callosum (Kawai et al., 2004).

Upon hearing a *wh*-interrogative at the beginning of a question, none of the left prefrontal ROIs showed significant neural engagement, and some left prefrontal ROIs, including the left aMFG and orbitofrontal regions, exhibited neural suppression (see 00:25 and 00:29 on **Video S5**). The lack of significant neural engagement upon hearing a *wh*-interrogative at the question beginning is consistent with the observations in our previous iEEG study (Iwaki et al., 2021). The current study revealed that the bilateral aMFG showed diminished inter-hemispheric connectivity via the anterior corpus callosum, while a suppressive neural information flow took place from right to left aMFG (see 00:09 on **Video S1** and 00:17 on **Video S2**). These findings may suggest an optimal allocation of neural resources (Norman and Bobrow, 1975) to optimize auditory processing, by reducing cognitive load in the prefrontal networks and by prioritizing the STG processing critical for analyzing speech sounds. The aMFG is implicated in cognitive control, such as selecting a required response while inhibiting automatic behavior (Frings et al., 2018). However, in the present study, patients were not assigned a dual task requiring cognitive control functions. Instead, their only option was to listen to and answer each of the questions. Thus, there was presumably no need to activate the cognitive control function located in the aMFG.

### 4.2. Common and distinct neural engagement patterns observed in naming questions that begin and end with a *wh*-interrogative

Common observations on neural engagement during tasks involving naming questions that begin and end with a *wh-*interrogative included enhanced left intra-hemispheric functional connectivity upon hearing a verb following either an adverb or an object. In trials that began with a *wh*-interrogative, extensive neural engagement was observed across the left frontal and temporal lobe neocortices upon hearing a verb (the third phrase) following either an adverb or object (the second phrase). This engagement was characterized by an enhancement of intra-hemispheric functional connectivity via the arcuate fasciculus, accompanied by bidirectional, facilitatory neural flows between these regions (see 00:13-00:14 on **Video S1** and 01:01-01:13 on **Video S2**). Similarly, in trials that concluded with a *wh*-interrogative, hearing a verb (the second phrase) after either an adverb or object (the first phrase) led to enhanced intra-hemispheric functional connectivity between the left temporal and frontal lobes through the arcuate fasciculus, along with bidirectional, facilitatory neural flows across these areas (00:37 on **Video S4**).

Conversely, we observed a notable difference in neural dynamics between questions that begin and end with a *wh-*interrogative, as discussed below. The spatial extent of left intra-hemispheric functional connectivity enhancement upon hearing the second phrase was significantly greater when patients encountered a combination of ‘adverb or object’ and ‘verb’, compared to a combination of ‘*wh*-interrogative’ and ‘adverb or object’ (**Figure 3**). The former combination can more readily elicit a semantically relevant response, whereas the latter requires hearing an additional concrete phrase before formulating a response. The timing of this enhancement in left intra-hemispheric functional connectivity aligns with contemporary models that propose bidirectional neural interactions between the temporal and frontal lobe neocortices through the arcuate fasciculus (Poeppel et al., 2012; Bornkessel-Schlesewsky et al., 2015; Rolls et al., 2022). Collective evidence suggests that such neural interactions are crucial for the semantic and syntactic processing needed to comprehend a spoken sentence and generate a semantically relevant response. For example, an fMRI study of healthy adults demonstrated hemodynamic activation of the left IFG, STG, and MTG by tasks involving syntactic and semantic processes for combining two spoken words (Humphries et al., 2007; Graessner et al., 2021; Schell et al., 2017). Studies of healthy adults using resting-state fMRI and DWI tractography indicated the existence of direct white matter pathways and bidirectional effective connectivity from the left STG and MTG to the left pIFG and pMFG (Milton et al., 2021; Rolls et al., 2022). A lesion-to-deficit study of 134 stroke survivors demonstrated that long-term speech production impairment was causally associated with severe damage to the left arcuate fasciculus connecting the left pIFG and temporal lobe neocortex rather than to the left pIFG alone (Gajardo-Vidal et al., 2021).

### 4.3. Distinct neural engagement patterns in naming questions beginning with *what*, *when,* and *where*

In trials initiated with a *wh*-interrogative, neural engagement in the left pIFG around the offset of a verb (the third phrase) was greater in questions asking *what* compared to *where*. The current study revealed that this preferential engagement in response to *what* questions was associated with enhanced functional connectivity with the left MTG and STG via the arcuate fasciculus, as illustrated in **Figure 4**. The increased involvement of the left arcuate fasciculus during this period can be attributed to the notion that internally generating a verbal response to *what* questions requires a higher cognitive load for lexical selection due to the broader range of potential responses, compared to responding to *where* questions. Our behavioral analysis demonstrated that the correct response rate for *what* questions was significantly lower compared to *when* and *where* questions. Lesion-to-deficit, fMRI, and iEEG studies indicate that the left pIFG plays a role in selecting the appropriate word from a wide range of possible correct answers (Schnur et al., 2009; Cho-Hisamoto et al., 2015; Riès et al., 2015).

### 4.4. Connectivity modulations prior to responses

Between the offset of three-phrase question to the onset of a patient’s response, neural engagement expanded to include the prefrontal, premotor, and precentral gyri within the left hemisphere. These structures showed increased functional connectivity and bidirectional, facilitatory neural information flows across the left frontal lobe (see 00:16-00:18 on **Video S1**; 01:21-01:29 on **Video S2**). For example, in trials initiated with a *wh-*interrogative, at 100-300 ms following the offset of the question, the left pIFG, aMFG, and precentral gyri exhibited bidirectional, facilitatory neural information flows through the frontal lobe’s u-fibers (01:21 on **Video S2**). Additionally, the left precentral and posterior superior frontal gyrus (pSFG) showed bidirectional, facilitatory information flows 100-300 ms prior to the response onset (01:29 on **Video S2**). The observed neural information flows between the left pIFG and the precentral gyrus may reflect the transfer of a mental representation of the verbal response to the primary motor area (Flinker et al., 2015; Nakai et al., 2017; Lu et al., 2018; Coletta et al., 2024); whereas the neural information flows between the left pSFG and the precentral gyrus are considered to be involved in the planning and initiation of overt responses (Krainik et al., 2003; Hertrich et al., 2016; Nakai et al., 2017).

### 4.5. Connectivity modulations during overt responses

As patients responded overtly to the questions, neural engagement was observed simultaneously in the bilateral precentral and postcentral gyri. This engagement involved enhanced inter-hemispheric functional connectivity and bidirectional, facilitatory information flows across the corpus callosum (00:19-00:21 on **Video S1**). These bidirectional flows likely facilitate synchronous and symmetric movements of the mouth and vocal structures, crucial for speech initiation (Hoptman and Davidson, 1994; Hervé et al., 2013). The critical role of the corpus callosum in speech initiation is underscored by reports of mutism immediately following corpus callosotomy (Sussman et al., 1983; Crutchfield et al., 1994).

During these overt responses, the STG also showed neural engagement, marked by enhancement in functional connectivity and bidirectional, facilitatory neural information flows through the posterior corpus callosum. Additionally, the left arcuate fasciculus, connecting the left precentral-postcentral gyri with the STG, exhibited increased intra-hemispheric functional connectivity and bidirectional neural flows. These connectivity modulations, involving the STG, are thought to support the monitoring of one’s own speech for semantic and prosodic accuracy and the internal integration of sound representations with oral movements during speech production (Zheng et al., 2010; Greenlee et al., 2013; Cogan et al., 2014).

### 4.6. IQ effects on neural engagement

In general, individuals with higher IQs possess more proficient verbal skills, including more extensive vocabularies and more effective communication. However, the mixed model analysis failed to show a significant association between IQ and naming behaviors, such as response accuracy or response time. Instead, the current study demonstrated that higher IQ correlated with reduced neural engagement in the left pMFG during overt responses. This reduction suggests a more efficient use of neural function within the left pMFG, which is presumably involved in processing heard sentences. Our previous iEEG studies highlighted that intense activation of the left pMFG occurs prior to responses in auditory descriptive naming tasks. In contrast, naming responses to pictures or environmental sounds, which do not require sentence processing, elicited much less activation in the left pMFG (Nakai et al., 2019; Kitazawa et al., 2023). This leads us to hypothesize that the left pMFG of individuals with higher IQs may complete sentence processing more rapidly during verbal responses. Conversely, the left pMFG of individuals with lower IQs might engage in sentence processing in a more prolonged manner while responding. Our observation is consistent with the results of a previous iEEG study of multilingual patients. The study demonstrated that the spatial extent of picture naming-related high-gamma augmentation was greater when they were required to respond in their second language compared to their first language, suggesting that processing a less proficient language may require additional cortical resources (Cervenka et al., 2011).

### 4.7. Novelty

This study pioneers a method by visualizing task-related modulations in both functional and effective connectivity via direct white matter pathways. Given the complexity and extensive volume of data, we chose animated videos over static images to comprehensively present our findings. The outcomes are consolidated into a video atlas that introduces a novel neurobiological framework, potentially enhancing understanding of human language processing in the brain.

We opted for a transfer entropy-based effective connectivity analysis, which quantifies unidirectional information flow by calculating one site’s ability to enhance the predictive capability about the future state of another (Ito et al., 2011). This choice was informed by our track record of applying this method to intracranial EEG data (Firestone et al., 2023). Analyzing trials that started with a *wh*-interrogative, we found that the facilitatory neural information flows were significantly stronger in the left hemisphere compared to the right when a verb was heard after an adverb or object. Furthermore, these information flows were bidirectional, primarily involving the left arcuate fasciculus. In the present study, our aim was to assess neural communications through the white matter directly connecting two ROIs. To ensure accuracy, the following criteria needed to be satisfied to confirm the occurrence of facilitatory neural information flows at any given moment:

[a] The two ROIs must be connected by a direct white matter tract, as evidenced by DWI tractography. This criterion aims to minimize the likelihood of erroneously identifying neural information flows between two cortical regions (e.g., between the right occipital and left frontal lobe cortical regions). Even if there is temporal coupling of neural engagement between the right occipital and left frontal lobe regions, it remains implausible for these cortical areas to communicate through a direct cortico-cortical white matter pathway. The current study focused on direct neural communications, explicitly excluding the visualization of multi-synaptic or indirect connectivity pathways.

[b] There must be a significant enhancement in functional connectivity through the white matter tract between the two ROIs. Specifically, the ROIs need to exhibit significant, simultaneous, and sustained (for at least 50 ms) high-gamma augmentation. In this study, the Type I error rate for detecting a functional connectivity enhancement between at least one pair of ROIs was found to be exceedingly low, at most 0.00000006. Consequently, the Type I error rate for identifying facilitatory neural information flows is similarly negligible.

### 4.8. Methodological considerations

Subsets of white matter pathways, despite showing significant functional connectivity enhancement, did not demonstrate facilitatory neural information flows as determined by our transfer entropy-based effective connectivity analysis. These findings warrant cautious interpretation. The Matlab script adapted from the algorithm described in Ito et al., 2011 required at least 20 time points to estimate the strength of transfer entropy-based effective connectivity. Consequently, in the present study, we quantified neural information flows using 200-ms time windows. Effective connectivity analysis, based on temporal changes in high-gamma amplitudes, implicates neural information flows when a high-gamma modulation in one region predicts subsequent modulation in another region. This computational method inherently requires a certain length of data points for reliable analysis. Previous iEEG studies have similarly inferred that effective connectivity assessments, such as those using Granger causality analysis, require data spanning hundreds of milliseconds (Korzeniewska et al., 2011; Johnson et al., 2018). Furthermore, video visualizations effectively interpret the direction of neural information flow when representing circles move continuously along white matter streamlines for a sustained duration. As a result, we could not achieve the same time resolution for effective connectivity analysis as we did for functional connectivity analysis. Due to these technical constraints, our effective connectivity analyses primarily aimed to clarify the direction of neural information flows when functional connectivity was enhanced during naming in response to questions beginning or ending with *wh-*interrogatives (**Videos S1-S4**). The possibility that the 200-ms analysis epoch, comprising twenty 10-ms time bins, was insufficient to capture neural information flows cannot be discounted. Yet, extending the analysis epoch might compromise the temporal resolution of the effective connectivity analysis. In contrast, functional connectivity could be quantified and visualized every 10 ms, provided that the connectivity modulations persisted for at least 50 ms.

Given that transfer entropy measures conditional mutual information in’bits’ (Schreiber 2000), we had to binarize the continuous high-gamma amplitude data to align with the requirements of our analysis equation. This binarization process might have led to a loss of sensitivity in detecting neural information flows across some pathways. Additionally, bivariate transfer entropy evaluates the information transfer from one time series at an ROI to another ROI, without considering potential influences from a third (or additional) ROI(s). Future studies should explore task-related effective connectivity using alternative algorithms, such as univariate or multivariate Granger causality analyses (e.g., Korzeniewska et al., 2011; Flinker et al., 2015; Rocchi et al., 2021) and graph theory metrics (e.g., Meunier et al., 2020; Yellapantula et al., 2021). Replicating similar patterns using distinct algorithms would provide additional credibility to our findings on pairwise effective connectivity.

We defined the presence of effective connectivity between two regions only when the following criteria were met: [1] a direct white matter streamline, [2] statistically-significant functional connectivity, and [3] transfer entropy-based neural information flows. This stringent criterion was applied to ensure that all visualized neural information flows are biologically plausible. While this approach enhances the plausibility of the visualized connectivities, it is likely that some functional and effective connectivity pathways, although present, were not captured in this study. For instance, none of the study patients had intracranial EEG sampling from thalamic nuclei, preventing the delineation of indirect connectivity mediated by the thalamus (Stitt et al., 2018).

The present study aimed to assess network dynamics by analyzing temporal changes in high-gamma modulations between two ROIs directly connected via white matter streamlines. This analytic approach partly relies on the notion that inter-regional communication occurs through high-frequency mechanisms. Our previous iEEG study identified auditory naming-related high-gamma co-augmentation at distant brain regions in individual patients, with these regions exhibiting intense effective connectivity, as defined by cortical responses induced by single-pulse electrical stimulation (Sonoda et al., 2021). In the present study, lower spectral frequency bands below high-gamma were not evaluated. Therefore, the absence of high gamma-based functional connectivity enhancement in a pathway at a specific moment does not necessarily exclude the presence of functional connectivity at lower spectral frequencies. For example, a study investigating visual memory processes using iEEG data from 294 patients with drug-resistant focal epilepsy found that connectivity networks assessed through task-related augmentation of broadband gamma power (30–100 Hz) were desynchronized across distant brain regions during memory encoding and retrieval (Solomon et al., 2017). In contrast, connectivity networks assessed using theta activity (3–8 Hz) were often synchronized across brain regions, with the extent of this synchrony positively correlating with broadband gamma power. Accordingly, the investigators suggested that low-frequency mechanisms may facilitate inter-regional communication (Solomon et al., 2017).

In the current study, no individual patient with bilateral recordings had electrode coverage of homotopic, non-epileptic gyri directly connected by the corpus callosum. Consequently, it is not feasible to apply our connectivity analysis methods to quantify interhemispheric neural interactions at the individual patient level. Furthermore, individual analyses with limited electrode coverage would be underpowered; each patient would need at least several hundred naming trials to achieve the same signal-to-noise ratio that a group-level analysis provides, and such a design would be infeasible in a clinical setting. To address the challenge of limited spatial sampling, investigators have integrated iEEG data from multiple patients into a standard brain to construct a single “Virtual Brain” and infer neural interactions within that framework (e.g., Kunii et al., 2013a; Burke et al., 2014; Avanzini et al., 2016; Solomon et al., 2017). While none of these studies employed transfer entropy-based effective connectivity analysis as used in the present study, the virtual brain framework enables systematic estimation of the timing of high-gamma amplitude modulations across multiple sites and interpretation of neural interactions using group summary data. It is reasonable to expect that the temporal dynamics of task-related high-gamma amplitude modulations in specific ROIs would be qualitatively consistent across individuals (Nakai et al., 2017; 2019). Furthermore, task-related high-gamma augmentation is characterized by a broadband increase in amplitude that is not confined to a patient-specific narrow frequency band (Lachaux et al., 2012). We acknowledge that it is unrealistic to expect the phase of high-gamma activity in ROIs to align across individuals. Consequently, in this study, we did not employ connectivity analyses reliant on coherence or traveling waves which depend on the phase of iEEG waveforms. Such analyses are inherently suitable only for individual-level investigations.

Neuroscientists commonly use standard brain templates, assuming that they are accurate representations of an average patient brain. In our study, we operated under a similar premise, positing that both patients and healthy individuals possess comparable white matter streamlines. The approach of combining open-source DWI data with individual iEEG data to investigate human brain network dynamics has received validation from two independent research groups (Mitsuhashi et al., 2021; Sonoda et al., 2021; Azeem et al., 2023; 2024). Notably, a previous iEEG study of ours successfully demonstrated the viability of a multimodal analysis by integrating iEEG measurements from non-epileptic cortical areas of epilepsy patients with DWI data from healthy individuals. We found that the lengths of white matter streamlines between language areas measured in individual tractography data were similar (Pearson’s r=0.8) to those measured using tractography data from 1,065 healthy individuals recruited through the Human Connectome Project (Sonoda et al., 2021).

While it is understood that white matter development undergoes significant changes in early childhood and more subtle changes during adolescence (Asato et al., 2010; Baum et al., 2022), the current study did not seek to explore age-related variations in DWI measures or to pinpoint abnormal DWI streamlines in patients with focal epilepsy. Rather, our objective was to delineate modulations in functional and effective connectivity through major white matter pathways that are consistently present across the general population. Considering that human head size achieves 95% of its adult dimensions by age 6 (Bastir et al., 2006), our methodology aligned with that of prior studies (Kitazawa et al., 2023; Ueda et al., 2024) in excluding patients with significant brain malformations affecting key brain sulci. Moreover, we excluded brain regions influenced by epileptiform discharges or structural lesions from our iEEG-based connectivity analyses.

In the present study, we did not perform a direct statistical comparison of local high-gamma or connectivity measures associated with the processing of a *wh-*interrogative between the beginning and end of sentences. A *wh*-interrogative at the beginning of sentences is unpredictable, whereas at the end of sentences, it can be more readily anticipated. Furthermore, the neural processing of a *wh-*interrogative presented at the end, but not at the beginning of sentences, is expected to include syntactic analysis.

Considering the inter-individual variability in response times, we must cautiously interpret our connectivity data. It is reasonable to consider that the reliability of connectivity measures might decrease relative to the time elapsed between the analysis epoch and either the stimulus sound or the patients’ responses. For instance, expecting uniform cortical activity across all patients 1,000 ms after the third phrase offset is unrealistic. Some patients began to verbalize an answer 1,000 ms after the third phrase offset, while others may have still been retrieving an appropriate word during the same period. Therefore, we did not include such a delayed period in the analysis. Instead, we limited the connectivity analyses to 300 ms after the third phrase offset, when no patients had verbalized answers yet. Conversely, it is reasonable to assume that auditory perception occurs during stimulus phrase presentation and motor preparation occurs immediately before verbal responses. Thus, we assessed the neural dynamics supporting overt responses using iEEG signals time-locked to the response onset. Time-locking iEEG analysis to both stimulus and response is a standard practice to overcome inter-individual differences in response times. In the current study, inter-individual variability in response time cannot directly account for our observation that left-hemispheric functional connectivity enhancement was more extensive in questions ending rather than beginning with a *wh-*interrogative (**Figure 3B**), because the significant difference was noted upon hearing the second phrase. Likewise, variability in response time cannot directly account for our observation of more intense left pIFG high-gamma augmentation associated with questions beginning with *what* compared to *where*, because the significant difference was noted within 100 ms after the third phrase offset (**Figure 4A**).

In the ancillary analysis, we observed interhemispheric facilitatory neural information flows involving the STG, precentral, and postcentral gyri around response onset (**Videos S2 and S4**). As summarized in **Tables S2 and S3**, iEEG signals were available from 16 and 12 patients for the left and right STG, respectively; 14 and 17 patients for the left and right precentral gyrus, respectively; and 16 and 17 patients for the left and right postcentral gyrus, respectively. Using Spearman’s correlation tests, we confirmed that mean high-gamma dynamics across patients at each ROI remained highly consistent, even after excluding data from individual patients. Specifically, Spearman’s correlation (rho) values indicated that the mean high-gamma dynamics from 500 ms before to 300 ms after response onset remained stable (rho > +0.98) after excluding any single patient (**Table S3**). Furthermore, transfer entropy analyses consistently demonstrated interhemispheric facilitatory neural information flows between homotopic ROIs, even after removing patients with the shortest or longest response times (**Table S4**). Collectively, these analyses support the interpretation that the observed interhemispheric neural interactions preceding responses are unlikely to be substantially confounded by inter-patient variability in response timing. However, we cannot entirely exclude the possibility that failure to detect interhemispheric neural information flows between other ROIs with small patient numbers might reflect noisy high-gamma signals from individual patients.

This study is not designed to quantify or visualize the local, short-range connectivity within a given ROI. The delineation and validation of short-range U-fiber streamlines with tractography face significant challenges, including the issue of crossing fibers and the absence of a universally accepted gold standard for their identification (Wang et al., 2012; Movahedian Attar et al., 2020). Therefore, we have proceeded with the assumption that sites within the same gyrus exhibiting high-gamma co-augmentation (e.g., in the STG upon hearing phrases) are likely interconnected by local, short-range white matter pathways (Chang et al., 2010; Trumpis et al., 2021; Bhaya-Grossman and Chang, 2022; Norman-Haignere et al., 2022; Leonard et al., 2024).

## 5. CONCLUSION

We visualized functional and effective connectivity through specific white matter as Japanese speakers responded to auditory descriptive questions. Our findings suggest that neural interactions within left-hemispheric white matter may be enhanced during the processing of verbs associated with concrete objects or adverbs. The phrase order of auditory descriptive questions, even when the overall meaning remains similar, may affect the network dynamics in listeners.

## Abbreviations

aMFG: anterior middle frontal gyrus
ANOVA: analysis of variance
CI: confidence interval
CT: computed tomography
DF: degree of freedom
DWI: diffusion-weighted imaging
fMRI: functional magnetic resonance imaging
IQR: interquartile range
iEEG: intracranial electroencephalography
IQ: intelligence quotient
ITG: inferior temporal gyrus
MNI: Montreal Neurological Institute
MRI: magnetic resonance imaging
MTG: middle temporal gyrus
pIFG: posterior inferior frontal gyrus
pMFG: posterior middle frontal gyrus
pSFG: posterior superior frontal gyrus
ROI: region of interest
SOZ: seizure onset zone
SPG: superior parietal gyrus
STG: superior temporal gyrus.

## Acknowledgments

This work was supported by Intramural Research Grant 28-4 Clinical Research for Diagnostic and Therapeutic Innovations in Developmental Disorders (to M.I.), JSPS KAKENHI Grant Number 19K09494 (to M.I.), JSPS KAKENHI Grant Number JP22KJ0323 (to N.K.), JSPS Overseas Research Fellowships 202460451 (to K. Sakakura), MEXT Grant-in-Aid for Scientific Research on Innovative Areas 19H04890 (to K. Suzuki), Grant-in-Aid for Transformative Research Areas 20H05956 (to K. Suzuki), Grant-in-Aid for Scientific Research (B) 21H03779 (to K. Suzuki), and the National Institute of Health (R01NS064033 to E.A.; R01NS089659 to J.W.J.; 1F30NS129239 to E.F.).

---

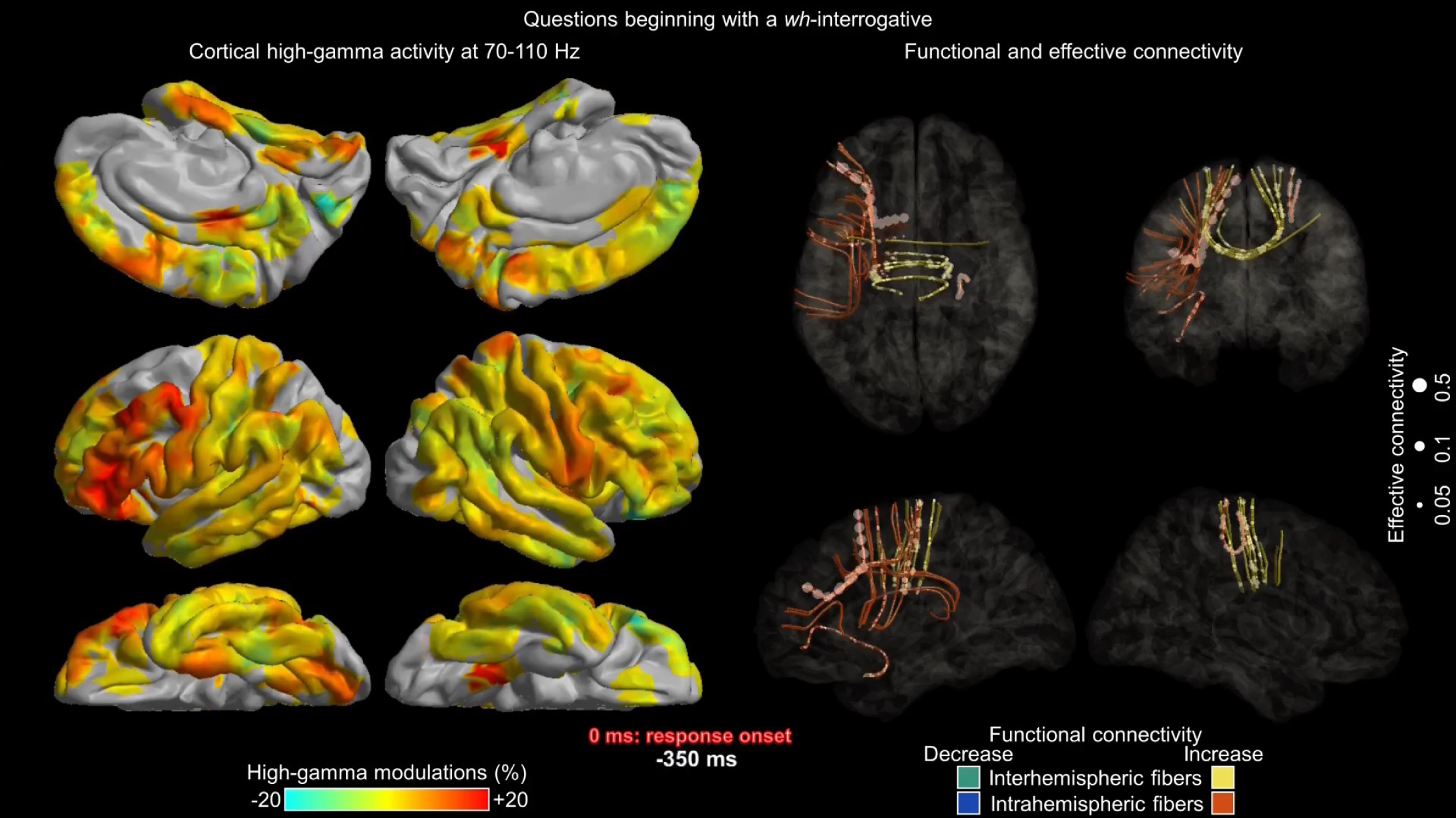

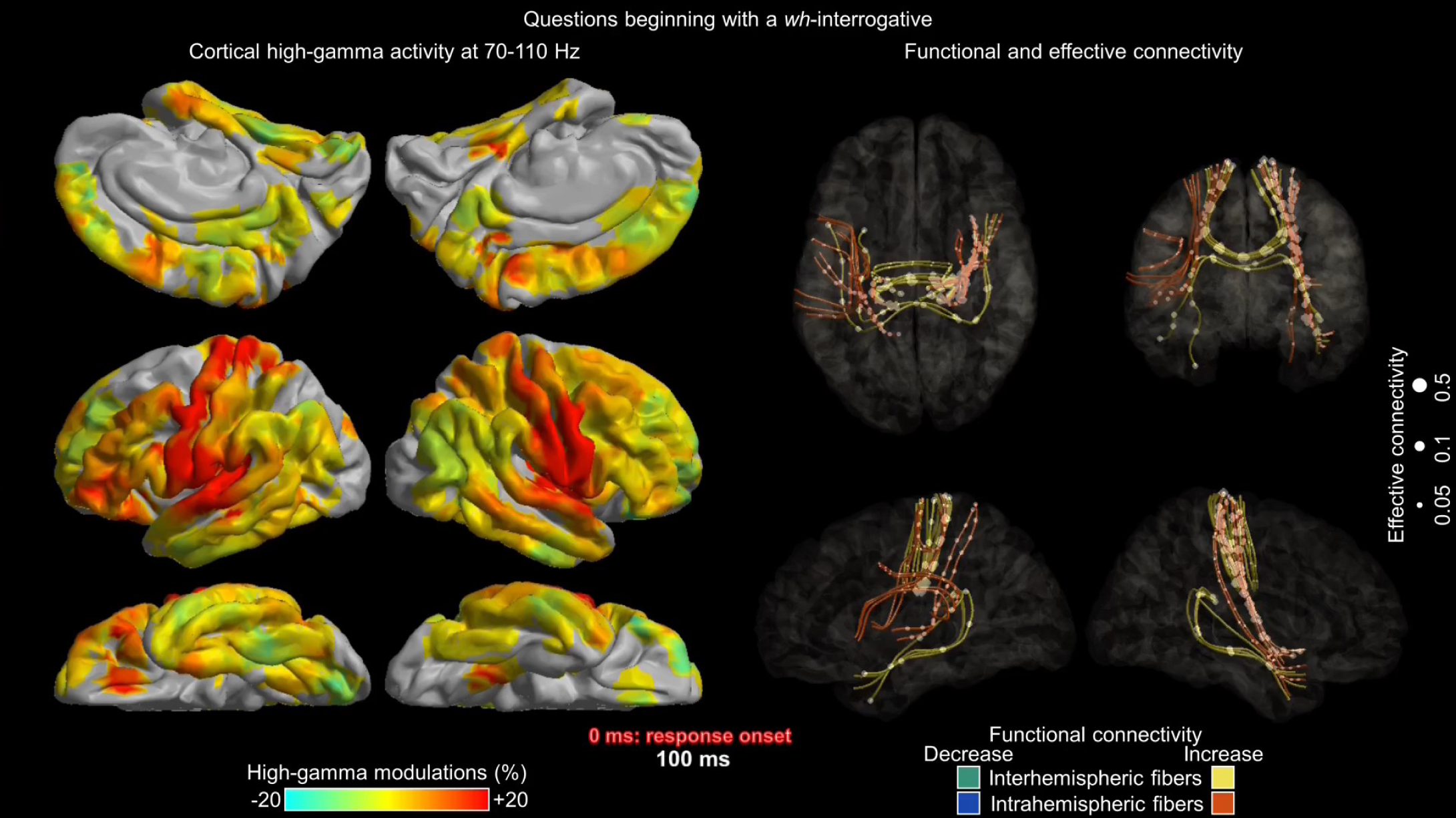

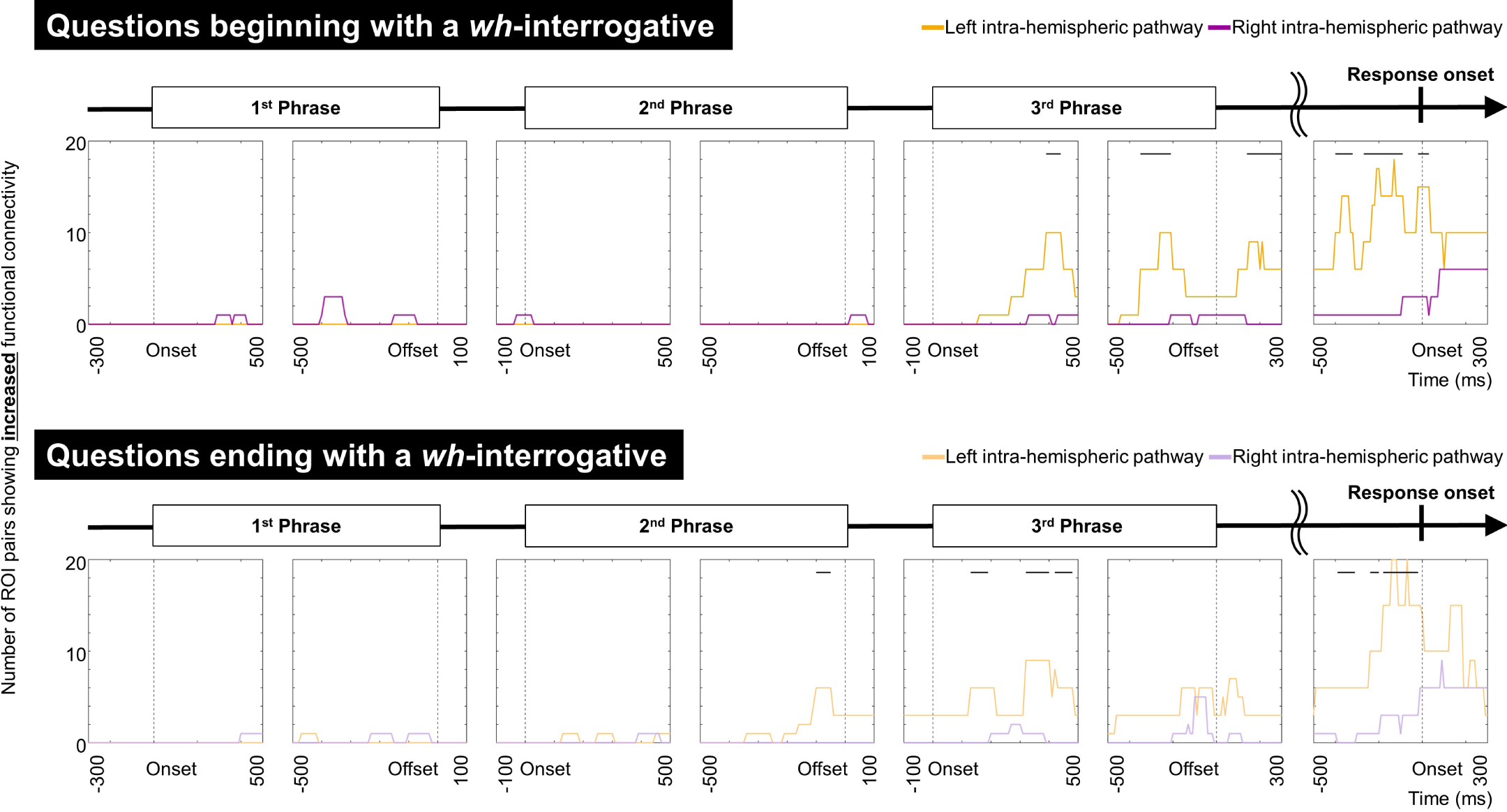

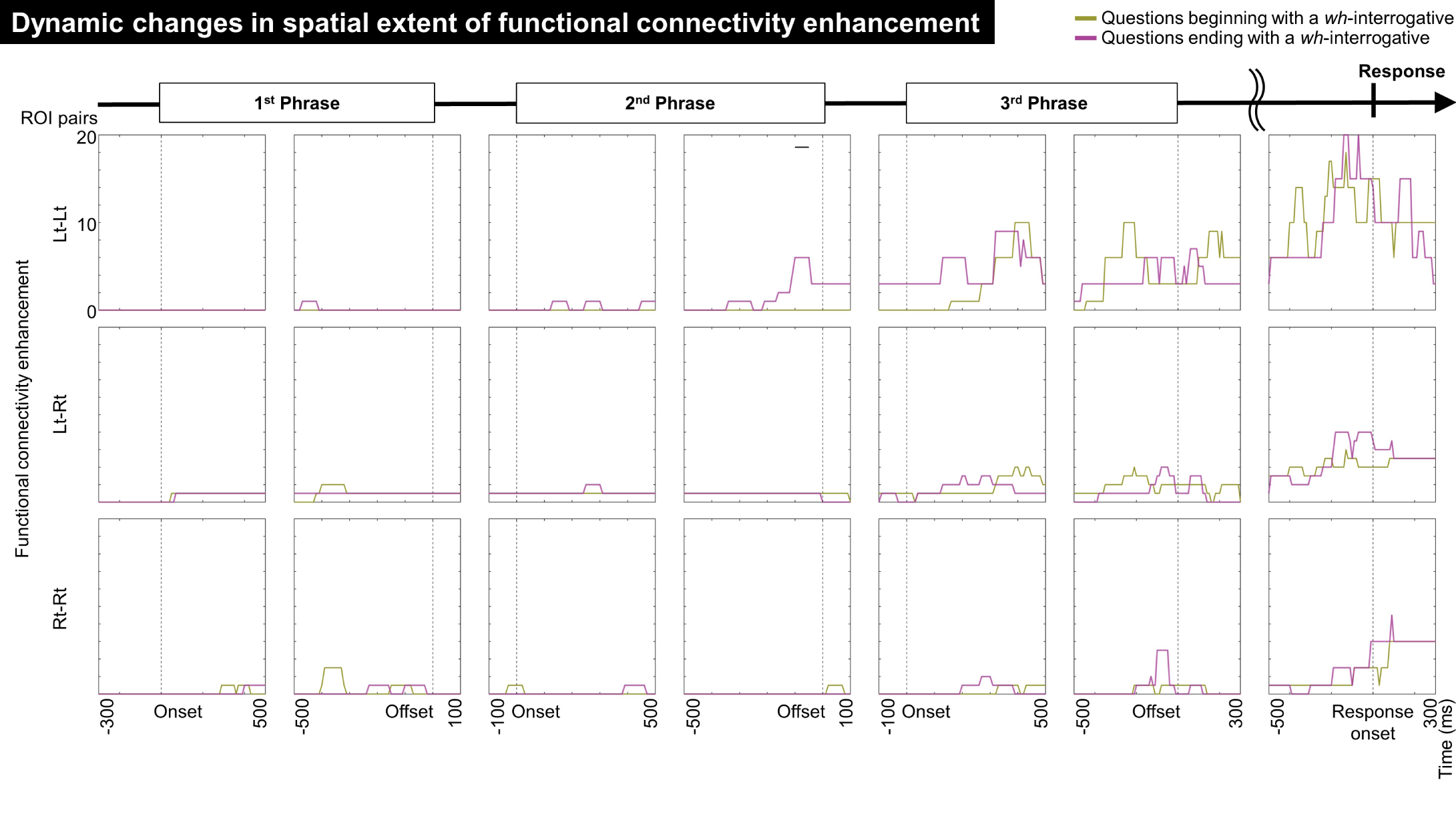

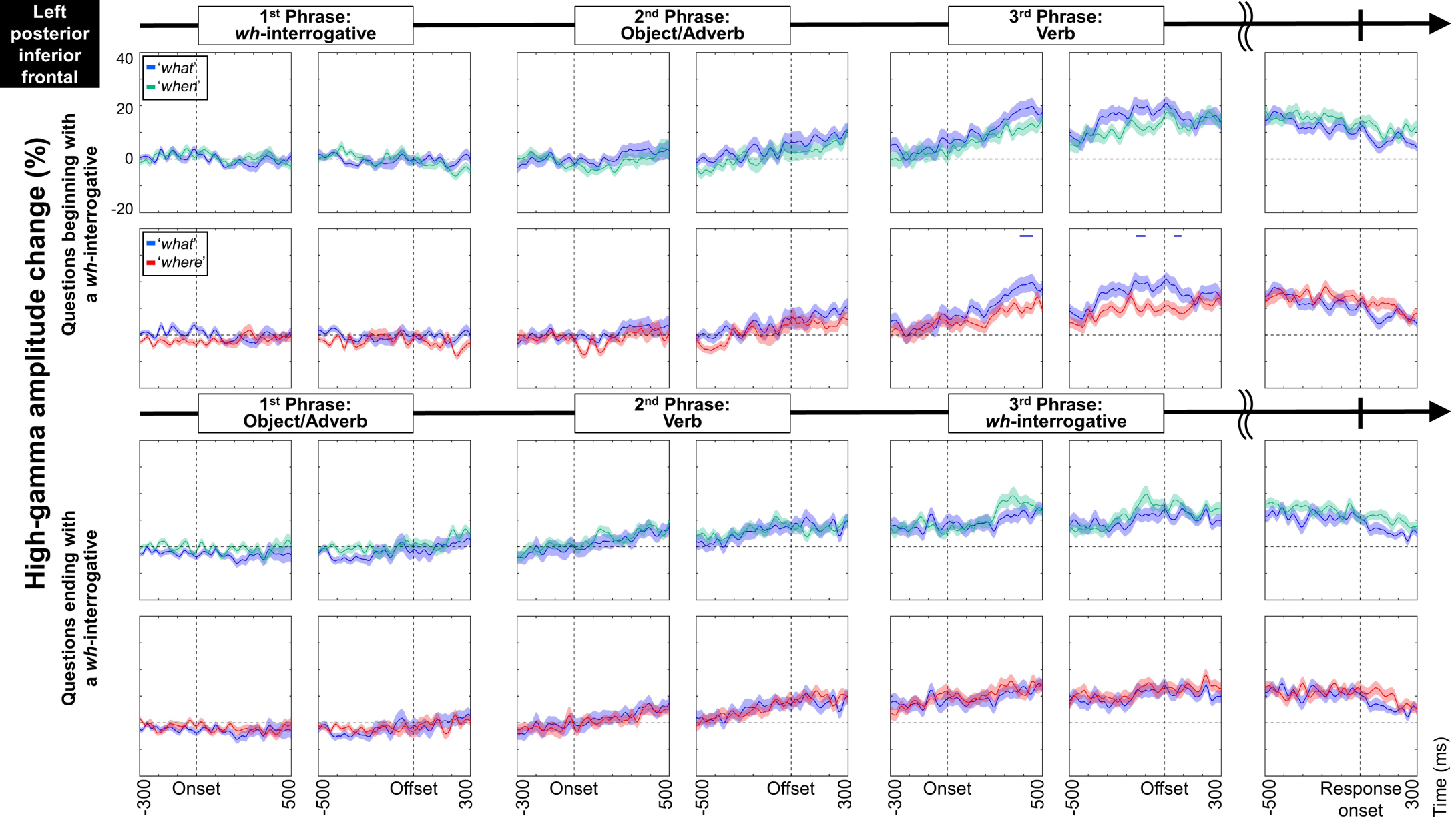

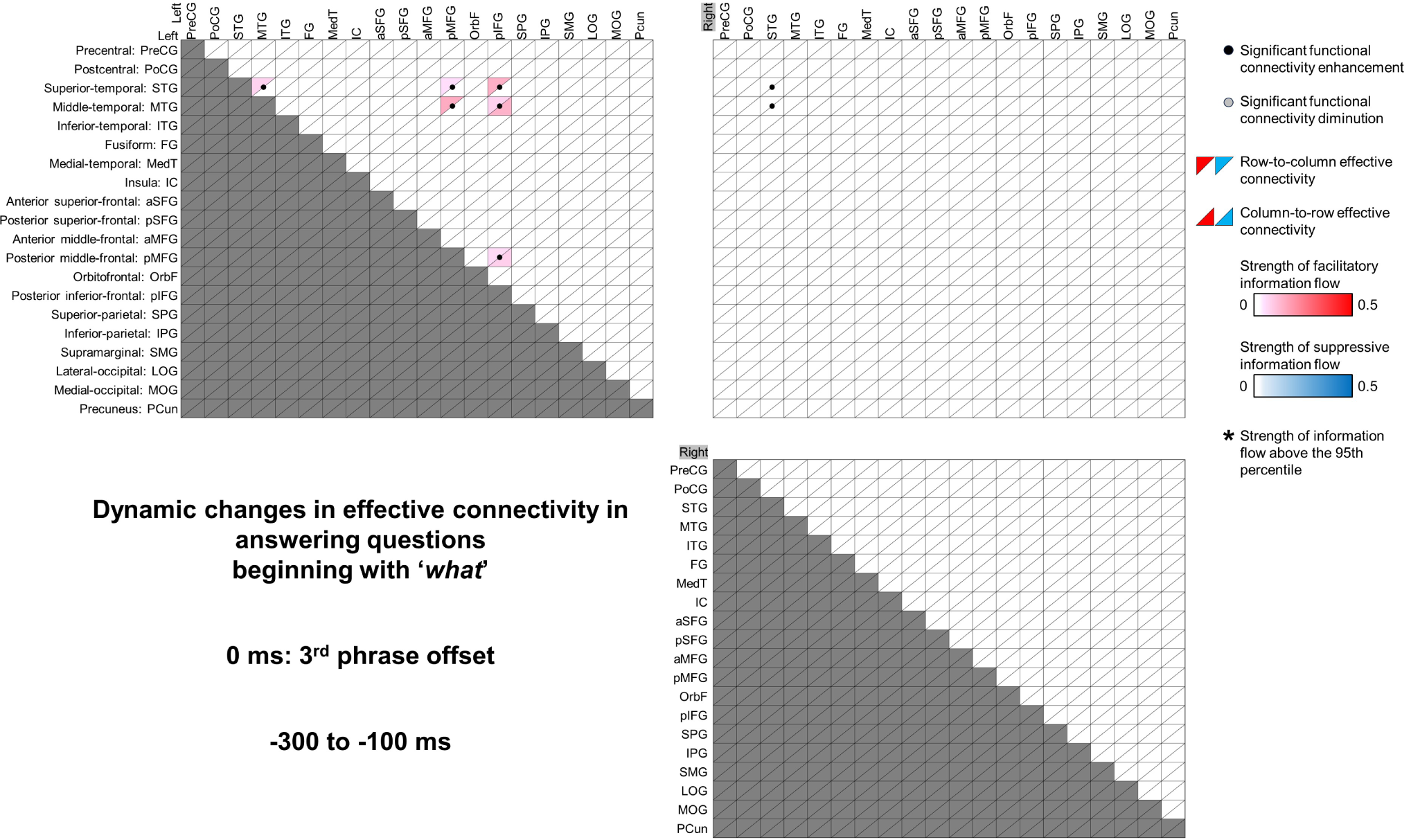

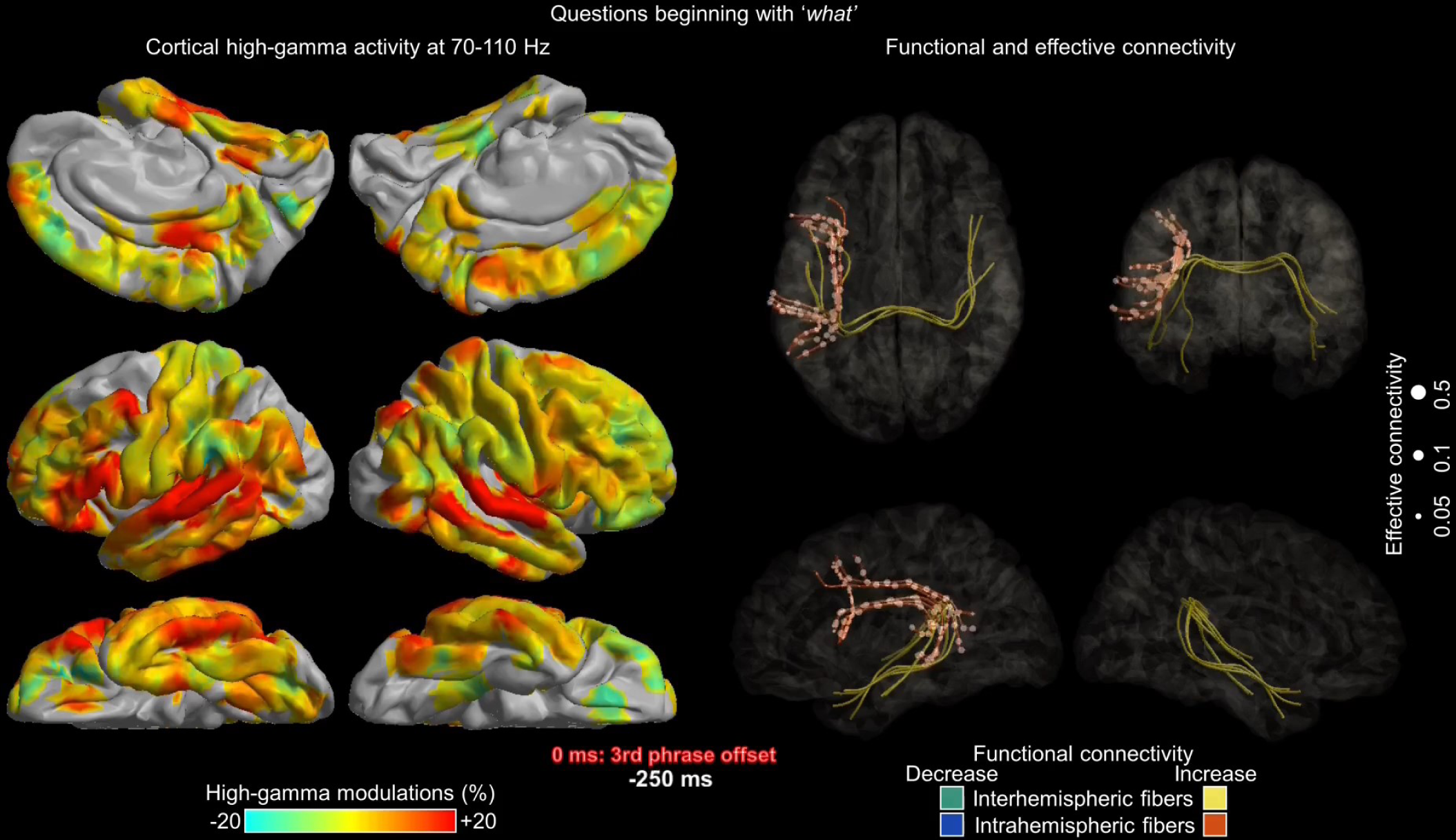

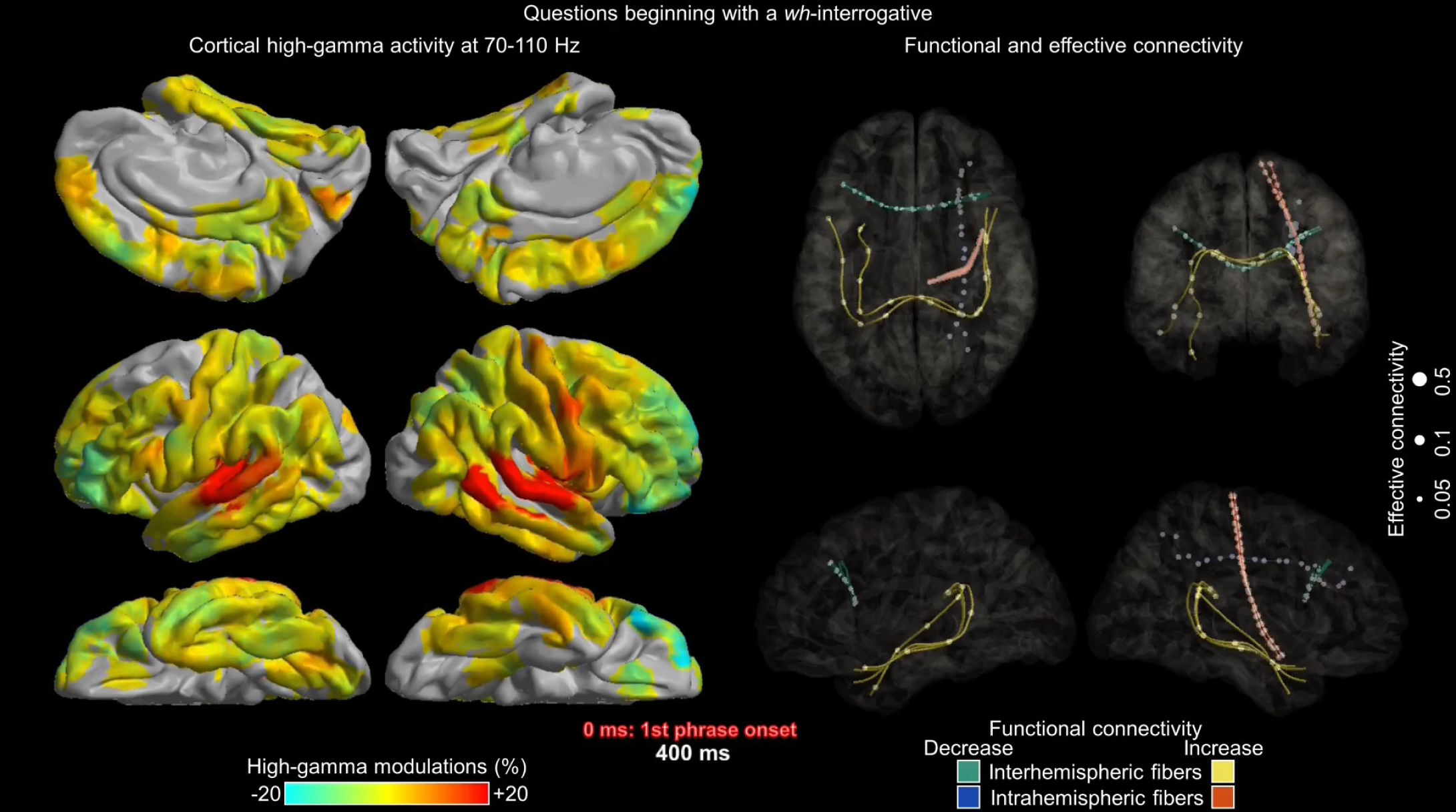

