## Supplementary tables, figure, and video legends for "Visualization of functional and effective connectivity underlying auditory descriptive naming"

This supplementary document includes:

**Tables S1-S4;**

**Figure S1;**

**Legends for Videos S1-S7.**

| **Patient** | **Age at surgery (years)** | **Sex** | **Sampled hemisphere** | **Seizure onset zone** | **Number of antiseizure medications** | **Full- scale IQ** | **Median response time (seconds)** | **Correct response rate** |
| --- | --- | --- | --- | --- | --- | --- | --- | --- |
| 1 | 26 | M | Lt | Lt T | 4 (CZP; LCM; TPM; VPA) | 96 | 1.84 | 0.99 |
| 2 | 33 | M | Lt | Lt TP | 3 (CBZ; CZP; LEV) | 98 | 2.32 | 0.99 |
| 3 | 36 | F | Lt | Lt T | 1 (LEV) | 104 | 1.10 | 0.99 |
| 4 | 32 | M | Rt | Rt FP | 2 (LCM; VPA) | 77 | 1.73 | 0.96 |
| 5 | 19 | F | Rt | Rt T | 5 (CBZ; CLB; DZP; PHT; VPA) | 74 | 1.44 | 0.74 |
| 6 | 54 | F | Rt | Rt FTP | 3 (CBZ; DZP; LEV) | 68 | 2.87 | 0.95 |
| 7 | 25 | M | Rt | Rt TP | 2 (LEV; ZNS) | 80 | 1.88 | 0.99 |
| 8 | 36 | F | Lt | Lt FT | 2 (LTG; PER) | 74 | 3.49 | 0.85 |
| 9 | 20 | M | Lt | Lt F | 2 (LTG; ZNS) | 50 | 1.25 | 0.92 |
| 10 | 16 | M | Lt | Lt T | 4 (CLB; CBZ; LEV; LTG) | 81 | 2.20 | 1.00 |
| 11 | 22 | M | Lt and Rt | Lt F | 4 (LTG; PER; VPA; ZNS) | 84 | 5.36 | 0.82 |
| 12 | 22 | M | Rt | Rt T | 2 (CBZ; LCM) | 83 | 0.84 | 0.93 |
| 13 | 25 | M | Lt | Lt T | 3 (LCM; LTG; PHT) | 68 | 0.84 | 0.94 |
| 14 | 10 | M | Lt | Gen | 4 (CBZ; CZP; LCM; VPA) | 58 | 1.45 | 0.96 |
| 15 | 6 | F | Lt | Lt C | 1 (LTG) | 77 | 3.79 | 0.74 |
| 16 | 17 | M | Lt | Lt O | 3 (CBZ; LCM; PER) | 67 | 0.60 | 0.72 |
| 17 | 26 | M | Lt and Rt | Rt T | 3 (CBZ; LCM; VPA) | 72 | 0.83 | 0.97 |
| 18 | 29 | F | Rt | Rt T | 3 (CLB; LTG; PER) | 114 | 4.09 | 0.91 |
| 19 | 15 | M | Lt and Rt | Lt F | 2 (DZP; LCM) | 91 | 0.82 | 0.95 |
| 20 | 37 | M | Lt and Rt | Rt O | 5 (CBZ; GBP; LEV; LTG; ZNS) | 63 | 2.20 | 0.95 |
| 21 | 38 | M | Lt | Lt T | 3 ( LEV; LTG; TPM) | 81 | 2.12 | 0.91 |
| 22 | 24 | F | Rt | Rt PO | 3 (CBZ; CLB; LTG) | 87 | 0.64 | 1.00 |
| 23 | 16 | F | Lt | Lt T | 2 (LEV; LCM) | 102 | 1.07 | 1.00 |
| 24 | 51 | F | Rt | Rt F | 4 (CBZ; CLB; LEV; LTG) | 84 | 1.13 | 0.89 |
| 25 | 23 | F | Lt | Lt F | 3 (LCM; LTG; VPA) | 76 | 0.57 | 0.97 |
| 26 | 11 | M | Rt | Rt O | 5 (CLB; LCM; LTG; PER; VPA) | 73 | 1.33 | 0.82 |
| 27 | 26 | F | Rt | Rt O | 3 (CLB; LCM; LTG) | 83 | 0.65 | 0.99 |
| 28 | 19 | M | Lt and Rt | Rt T | 4 (LCM; PER; PHT; VPA) | 83 | 1.55 | 0.89 |
| 29 | 19 | M | Lt and Rt | Lt T | 4 (CLB; ESM; LCM; TPM) | 58 | 2.19 | 0.91 |
| 30 | 27 | F | Rt | Rt F | 3 (CBZ; CLB; LEV) | 91 | 1.24 | 0.99 |
| 31 | 20 | M | Lt | Lt T | 5 (CLB; LEV; LTG; TPM; VPA) | 72 | 1.22 | 0.96 |
| 32 | 30 | F | Rt | Rt T | 3 (LCM; LTG; ZNS) | 88 | 2.16 | 0.92 |
| 33 | 17 | F | Rt | Rt F | 2 (LCM; LEV) | 74 | 2.50 | 0.79 |
| 34 | 21 | F | Lt | Lt TPO | 3 (LCM; LEV; LTG) | 70 | 1.21 | 0.97 |
| 35 | 11 | M | Lt | Lt T | 3 (LCM; LEV; LTG) | 59 | 1.72 | 0.77 |
| 36 | 33 | F | Rt | Rt F | 3 (CBZ; LEV; LTG) | 74 | 1.48 | 0.94 |
| 37 | 9 | M | Lt | Lt T | 5 (CLB; LEV; LTG; VPA; ZNS) | 83 | 2.69 | 0.78 |
| 38 | 37 | M | Lt and Rt | Rt T | 2 (LCM; LEV) | 83 | 2.62 | 0.93 |
| 39 | 22 | M | Lt | Lt FT | 3 (CBZ; PER; ZNS) | 95 | 1.60 | 0.93 |
| 40 | 40 | M | Lt | Lt T | 3 (LCM; LEV; LTG) | 88 | 1.76 | 1.00 |

**Table S1: Patient profile.** F: Female. M: Male. Lt: Left. Rt: Right. C: Central. F: Frontal. Gen: Generalized. T: Temporal. O: Occipital. P: Parietal. CBZ: Carbamazepine. CLB: Clobazam. CZP: Clonazepam. DZP: Diazepam. ESM: Ethosuximide. GBP: Gabapentin. LCM: Lacosamide. LEV: Levetiracetam. LTG: Lamotrigine. PER: Perampanel. PHT: Phenytoin. TPM: Topiramate. VPA: Valproic acid. ZNS: Zonisamide. The Kruskal-Wallis test showed no significant difference in age (p-value: 0.32) or response time (p-value: 0.75) across patients undergoing left-hemispheric, right-hemispheric, and bilateral iEEG sampling.

| **Region of interests** | **Left** | **Right** |
| --- | --- | --- |
| PreCG: Precentral | 67 (14) | 79 (17) |
| PoCG: Postcentral | 67 (16) | 69 (17) |
| STG: Superior-temporal (superior temporal and transverse temporal) | 90 (16) | 49 (12) |
| MTG: Middle-temporal | 54 (12) | 66 (15) |
| ITG: Inferior-temporal | 46 (10) | 29 (9) |
| FG: Fusiform | 36 (12) | 14 (5) |
| MedT: Medial-temporal (entorhinal and parahippocampal) | 9 (5) | 7 (4) |
| IC: Insula | 9 (4) | 28 (6) |
| aSFG: Anterior superior-frontal | 29 (3) | 49 (8) |
| pSFG: Posterior superior-frontal | 14 (3) | 19 (7) |
| aMFG: Anterior middle-frontal | 56 (10) | 78 (10) |
| pMFG: Posterior middle-frontal | 34 (6) | 39 (12) |
| OrbF: Orbitofrontal (BA 11, 12, and 47) | 19 (6) | 18 (7) |
| pIFG: Posterior inferior-frontal (BA 44 and 45) | 47 (12) | 57 (10) |
| SPG: Superior-parietal | 16 (8) | 34 (10) |
| IPG: Inferior-parietal | 30 (11) | 51 (12) |
| SMG: Supramarginal | 54 (15) | 82 (16) |
| LOG: Lateral-occipital | 8 (3) | 9 (5) |
| MOG: Medial-occipital (lingual and cuneus) | 27 (9) | 5 (3) |
| Pcun: Precuneus | 10 (4) | 15 (5) |
| pCG: Posterior cingulate (posterior cingulate and isthmus cingulate) | 12 (5) | 4 (3) |
| aCG: Anterior cingulate (rostral anterior cingulate and caudal anterior cingulate) | 2 (2) | 7 (4) |
| PCL: Paracentral | 7 (2) | 10 (2) |
| TP: Temporal pole | 4 (1) | 2 (1) |
| FP: Frontal pole | 0 (0) | 2 (1) |
| Total | 747 (26) | 822 (21) |

**Table S2: The exact number of artifact-free, nonepileptic electrode sites in given regions of interest (ROI).** Number in a parenthesis: number of contributing patients. The ROI-based analysis was performed using those with at least five cortical electrode sites derived from at least three patients in each hemisphere.

Contributing	Spearman’s rho value

ROI	patients	Mean	95%CI	P-value

Left STG	16	+0.996	+0.993 to +0.999	1.1 × 10^-35^

Right STG	12	+0.993	+0.989 to +0.998	4.6 × 10^-25^

Left PreCG	14	+0.990	+0.982 to +0.998	6.2 × 10^-26^

Right PreCG	17	+0.994	+0.990 to +0.997	4.1 × 10^-36^

Left PostCG	16	+0.990	+0.986 to +0.994	5.4 × 10^-33^

Right PostCG	17	+0.994	+0.992 to +0.996	3.8 × 10^-40^

**Table S3: Robustness of high-gamma dynamics at regions of interest (ROIs).** Using Spearman’s correlation, we determined whether the high-gamma amplitude dynamics between 500 ms pre-response onset and 300 ms post-response onset were consistent, even when a patient was excluded from the computation. For example, we repeated the Spearman’s correlation test 16 times at the left superior-temporal gyrus (STG). This analysis yielded the mean Spearman’s rho and its 95% confidence interval (95% CI). The one-sample t-test demonstrated that the mean Spearman’s rho value was greater than zero at all six ROIs showing interhemispheric, facilitatory neural information flow around response onset. High Spearman’s rho values suggest that high-gamma dynamics at each ROI are robust and were not driven by the effect of a single patient. PreCG: precentral gyrus. PostCG: postcentral gyrus.

Maximum neural information flow

Auditory	ROI	All patients	Patient with the	Patient with the

shortest response	longest response

stimulus	pairs	included	time excluded	time excluded

Rt→Lt STG	0.254	0.254	0.254

Beginning with	Lt→Rt PreCG	0.319	0.361	0.372

a *wh*-interrogative	Rt→Lt PreCG	0.144	0.302	0.302

Lt→Rt PostCG 0.216	0.216	0.216

Rt→Lt PostCG 0.429	0.429	0.429

Lt→Rt STG	0.483	0.483	0.483

Rt→Lt STG	0.104	0.104	0.104

Ending with	Lt→Rt PreCG	0.278	0.264	0.293

a *wh*-interrogative	Rt→Lt PreCG	0.293	0.464	0.340

Lt→Rt PostCG 0.446	0.446	0.446

Rt→Lt PostCG 0.164	0.237	0.164

**Table S4: Robustness of interhemispheric facilitatory information flows.** By computing transfer entropy, we determined whether the interhemispheric facilitatory information flows between 500 ms pre-response onset and 300 ms post-response onset were consistent, even when a patient with the shortest or longest response time was excluded from the computation.

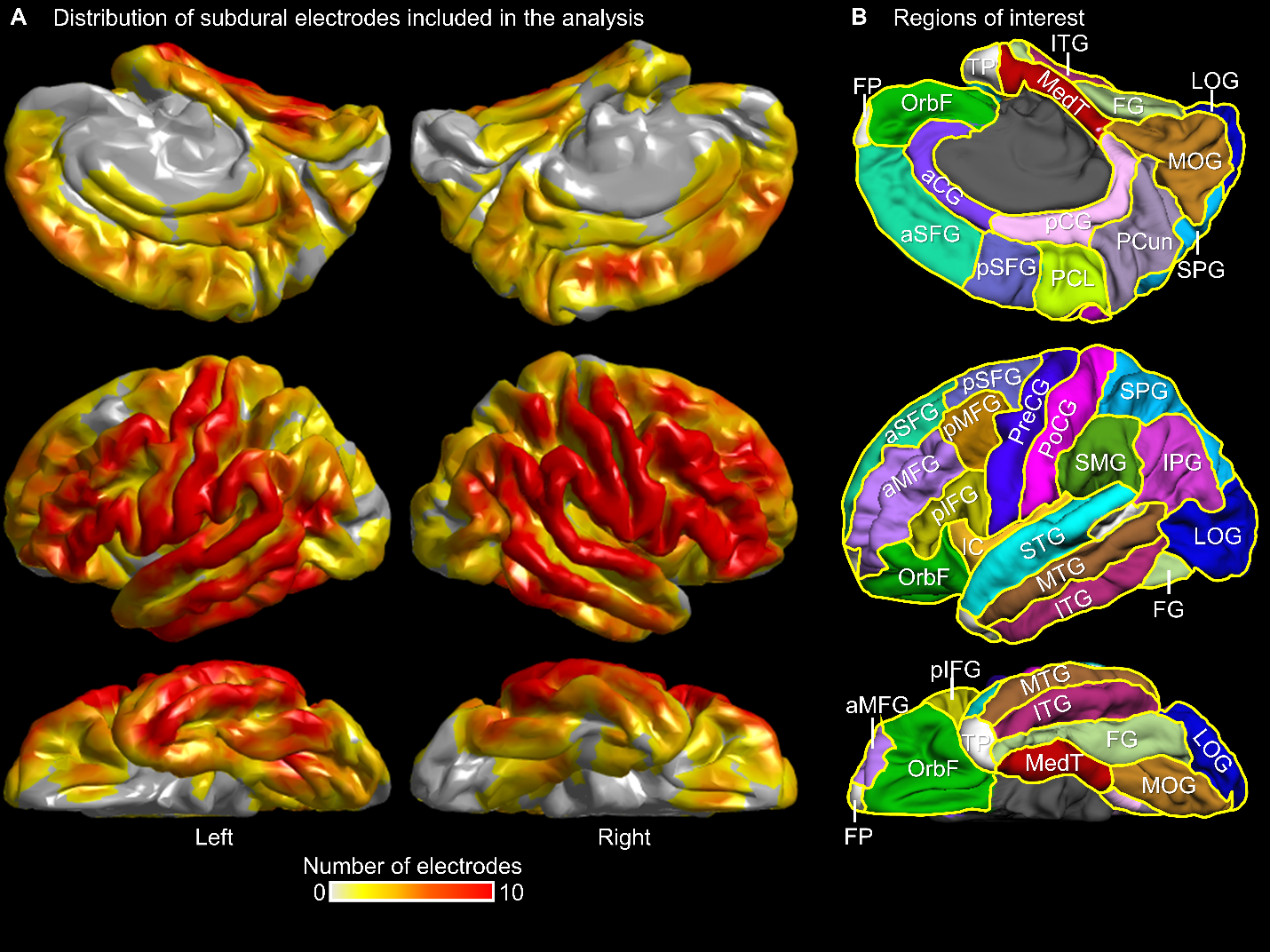

**Figure S1: Regions of interest (ROIs) and distribution of intracranial electrode sites included in the analysis.** (A) The distribution of artifact-free, nonepileptic electrode sites from the 40 patients. (B) ROI locations on the left hemisphere are presented. **Table S1** provides the full form of each abbreviation and the exact number of artifact-free, nonepileptic electrode sites in given ROIs.

**Legends for Videos S1-S7.**

**Video S1: Seven-dimensional connectivity atlas of naming responses to questions beginning with a *wh*-interrogative.** The video on the left demonstrates the dynamics of cortical high-gamma amplitude modulations, while the video on the right illustrates the dynamics of functional and effective connectivity during an auditory description naming task involving overt responses to three-phrase questions that begin with a *wh-*interrogative, followed by either an adverb or object, and then a verb. Yellow white matter streamlines indicate significant inter-hemispheric functional connectivity enhancement, with orange indicating significant intra-hemispheric enhancement. Conversely, green streamlines show significant inter-hemispheric connectivity diminution, and blue ones indicate intra-hemispheric diminution. The direction and strength of neural information flows are represented by the movement and size of white circles, respectively.

**Video S2: Dynamics of functional and effective connectivity during naming responses to questions beginning with a *wh*-interrogative.** This video illustrates the dynamics of functional and effective connectivity during naming responses to questions that begin with a *wh*-interrogative. Black circles represent enhanced functional connectivity between regions of interest (ROIs) via direct white matter pathways within a given 200-ms timeframe. Conversely, gray circles denote diminished functional connectivity. Colored triangles signify neural information flows from one ROI to another. Red triangles indicate facilitatory and blue ones indicate suppressive neural information flows. For instance, during a 200-ms period post the initial phrase's onset, a facilitatory neural information flow occurred from the left to the right superior temporal gyrus. An asterisk (*) highlights ROI pairs with neural information flow strengths exceeding the 95th percentile.

**Video S3: Seven-dimensional connectivity atlas of naming responses to questions ending with a *wh*-interrogative.** The video on the left demonstrates the dynamics of cortical high-gamma amplitude modulations, while the video on the right illustrates the dynamics of functional and effective connectivity during an auditory description naming task involving overt responses to three-phrase questions that begin with either an adverb or object, followed by a verb, and then a *wh-*interrogative. Yellow white matter streamlines indicate significant inter-hemispheric functional connectivity enhancement, with orange indicating significant intra-hemispheric enhancement. Conversely, green streamlines show significant inter-hemispheric connectivity diminution, and blue ones indicate intra-hemispheric diminution. The direction and strength of neural information flows are represented by the movement and size of white circles, respectively.

**Video S4: Dynamics of functional and effective connectivity during naming responses to questions ending with a *wh*-interrogative.** This video illustrates the dynamics of functional and effective connectivity during naming responses to questions that ending with a *wh*-interrogative. Black circles represent enhanced functional connectivity between regions of interest (ROIs) via direct white matter pathways within a given 200-ms timeframe. Conversely, gray circles denote diminished functional connectivity. Colored triangles signify neural information flows from one ROI to another. Red triangles indicate facilitatory and blue ones indicate suppressive neural information flows. For instance, during a 200-ms period post the initial phrase's onset, a facilitatory neural information flow occurred from the left to the right superior temporal gyrus. An asterisk (*) highlights ROI pairs with neural information flow strengths exceeding the 95th percentile.

**Video S5: Cortical high-gamma modulations during naming responses to questions beginning and ending with a *wh*-interrogative.** This video illustrates the dynamics of cortical high-gamma modulations during an auditory description naming task, presenting the mean high-gamma amplitude alongside its 99.99% confidence interval. Light blue indicates cortical high-gamma amplitude modulations when responding to questions that begin with a *wh*-interrogative, followed by either an adverb or object, and then a verb. Magenta represents modulations when the questions start with an adverb or object, are followed by a verb, and end with a *wh-*interrogative.

**Video S6: Functional connectivity dynamics during naming responses to various question types.** This video demonstrates the dynamic changes in the spatial extent of functional connectivity, represented by the number of ROI pairs, during the process of responding to different types of questions. The study analyzes both intra-hemispheric and inter-hemispheric ROI pairs, with a total of 190 intra-hemispheric ROI pairs in each hemisphere and 400 inter-hemispheric ROI pairs available. A horizontal bar within the video marks the time points at which the proportion of ROI pairs showing significant functional connectivity enhancement (or diminution) significantly differed between the different question types.

**Video S7: Cortical high-gamma modulations during naming responses to questions asking *what, when,* and *where*.** This video illustrates the dynamics of cortical high-gamma modulations during an auditory description naming task, presenting the mean high-gamma amplitude alongside its standard error shade. Blue line: questions beginning with *what*. Green lines: those beginning with *when*. Magenta line: those beginning with *where*. A horizontal bar indicates a time period where high-gamma amplitudes significantly differed between question types.
